# Disease-associated patterns of acetylation stabilize tau fibril formation

**DOI:** 10.1101/2023.01.10.523459

**Authors:** Li Li, Binh Nguyen, Vishruth Mullapudi, Lorena Saelices, Lukasz A. Joachimiak

## Abstract

Assembly of the microtubule-associated protein into tauopathy fibril conformations dictates the pathology of a diversity of diseases. Recent cryogenic Electron Microscopy (cryo-EM) structures have uncovered distinct fibril conformations in different tauopathies but it remains unknown how these structures fold from a single protein sequence. It has been proposed that post-translational modifications may drive tau assembly but no direct mechanism for how modifications drive assembly has emerged. Leveraging established aggregation-regulating tau fragments that are normally inert, we tested the effect of chemical modification of lysines with acetyl groups on tau fragment conversion into amyloid aggregates. We identify specific patterns of acetylation that flank amyloidogenic motifs on the tau fragments that drive rapid fibril assembly. To understand how this pattern of acetylation may drive assembly, we determined a 3.9 Å cryo-EM structure of an amyloid fibril assembled from an acetylated tau fragment. The structure uncovers how lysine acetylation patterns mediate gain-of-function interactions to promote amyloid assembly. Comparison of the structure to an ex vivo tau fibril conformation from Pick’s Disease reveals regions of high structural similarity. Finally, we show that our lysine-acetylated sequences exhibit fibril assembly activity in cell-based tau aggregation assays. Our data uncover the dual role of lysine residues in limiting aggregation while their acetylation leads to stabilizing pro-aggregation interactions. Design of tau sequence with specific acetylation patterns may lead to controllable tau aggregation to direct folding of tau into distinct folds.

## Introduction

The deposition of the microtubule-associated protein in beta-sheet-rich amyloid conformations is a hallmark of many neurodegenerative diseases. These diseases are classified as tauopathies and include Alzheimer’s (AD), Corticobasal degeneration (CBD), Pick’s (PiD), chronic traumatic encephalopathy and frontotemporal dementia with tau (FTD-tau)^1^. Recent ex vivo cryo-EM structures of tau fibrils isolated from tauopathy patient material have resolved a series of distinct structures (i.e. structural polymorphs) each linked to a different disease ^2–5^. Moreover, each structural polymorph of tau has been proposed to encode the capacity to replicate their shape in a prion-like manner, spread their conformation throughout the brain, and to cause progressive neurodegeneration^6–9^. Despite this strong link to disease, the intrinsically disordered protein tau under normal conditions is aggregation-resistant and its aggregation must be induced using polyanions or pathogenic seeds to form beta-sheet-rich amyloids^10–13^. It thus remains unknown how tau converts into pathogenic states capable of initiating conformation-specific tauopathies.

Short amyloid-forming motifs (^274^VQIINK^280^, ^306^VQIVYK^311^ and ^337^VEVKSE^342^) control tau assembly into fibrils by engaging in hetero-typic interactions with other parts of the sequence. The amyloid motifs in tau localize to the beginning of each of the four repeats (i.e. R1, R2, R3 and R4) in the repeat domain^1^. Each of the amyloid forming motifs is preceded by a P-G-G-G sequence motif that stabilizes beta-turn conformations^14^. These beta-turn motifs have been shown to regulate the aggregation capacity of tau by engaging side-chain interactions with the adjacent amyloid motifs^14–16^. We and others have shown that perturbation of beta-turn-based regulatory elements in a monomer can lead to the formation of a seed-competent monomer^12,13,16^. The seeding-competent monomer species can be isolated from AD patients and tauopathy mouse brains but are absent in control tissues^12,13,17^. These regulatory elements can be recapitulated with wildtype tau peptides that span the amyloid motif and the preceding sequence including the P-G-G-G motif in 4 repeat (4R) tau (i.e. R1R2, R2R3, R3R4 and R4R’) as well as 3 repeat (3R) tau in which the second repeat is spliced out creating a new junction between repeats 1 and 3 (i.e. R1R3). Disease-associated mutations in tau linked to FTD-tau localize to these regulatory elements and increase aggregation capacity into fibrils compared to wildtype ^18^. We hypothesized that dysregulation of these elements may preferentially expose different amyloid motifs enabling the combinatorics of their exposure to stabilize intermediates towards distinct structural polymorphs.

For many years, post-translational modifications (PTMs) such as phosphorylation have been implicated in conversion of tau into pathogenic conformations^19–22^. However, recent data have suggested that phosphorylation may not be the emergent perturbation that drives the formation of the first pathogenic tau species but rather accumulates once the assemblies are formed ^17^. Detection of phosphorylated tau is the *de facto* method for disease diagnosis in blood ^23^ but also for detection of tau pathology in post-mortem tissues^24,25^. Phosphorylation on tau is relatively ubiquitous but excluded from the repeat domain with the exception of AD-specific modifications on serine S262 that lie outside of the AD fibril core^17,26^. Consistent with this observation, phospho-modifications in the repeat domain inhibit tau assembly ^27^. Other common PTMs such as acetylation and ubiquitination have similarly been implicated in driving progression of disease through promoting aggregation but the mechanism remains unknown ^17,26^. A recent structural and proteomic analysis of AD and CBD fibrils proposed that acetylation and ubiquitination may be important for control of structural polymorph assembly ^28^. Additionally, others have shown that acetylation-specific antibodies can detect early pathogenic species^29^. It is not known how acetylation may influence tau aggregation. It is possible that acetylation on lysines directly influences aggregation by stabilizing pro-aggregation interactions, disrupting protective interactions or reducing interaction with microtubules^30–32^. Moreover, it is possible that acetylation may influence the process by blocking ubiquitination sites required for subsequent degradation. Sites of acetylation on tau have been reported throughout the entire sequence including in the repeat domain surrounding the amyloid motifs including lysines K267, K274, K280 and K281 in repeats 1 and 2 ^26^. Such sites have also been detected in repeats 3 and 4 at lysines K311 and K353^26^. The acetyl transferase enzymes involved in this process are not known but histone deacetylases (HDACs) and SIRT1 have been implicated in their removal. HDAC6 has been implicated as the deacetylase that may regulate removal of acetylation from lysines on tau ^30^. Specific modifications surrounding the amyloid motifs (i.e. K274, K280 and K311) have been proposed to alter Hsp70 recruitment thus reducing chaperone-mediated autophagy ^33^. Furthermore, tau encoding acetylated K280 generated using semi-synthetic methods increases heparin-induced aggregation propensity 3-fold compared to unmodified tau yielding a mixture of oligomers and short fibrils ^34^. Thus the proximity of disease-associated acetylation sites on tau to our established aggregation regulatory elements^14^ suggests that we may be able to test the role of lysine acetylation in mediating early conformational changes to drive tau assembly.

Here we show that disease-associated patterns of tau acetylation on minimal tau fragments encoding regulatory elements can drive amyloid assembly. We first use a chemical approach to modify tau fragments that span the R1R2, R2R3, R3R4 and R4R’ in 4R tau and R1R3 in 3R tau. We show that saturating chemical acetylation of R1R2, R2R3 and R1R3 leads to rapid fibrillar assembly while the unmodified sequences remain inert. We next use synthetic sequences with site specific acetylation to uncover patterns of modification that regulate the assembly of tau fragments. We also test commonly used lysine to glutamine substitutions and show that similar modifications can lead to their assembly with drastically different fibril morphologies. We use cryo-EM to determine a fibril structure of one of our R1R2 acetylated peptides encoding disease-associated acetylation sites at residues K274 and K280. The structure uncovers interactions mediated by acetylation that balance hydrogen bonding and nonpolar contacts to stabilize the structure not possible by a lysine side chain. The structure also revealed that specific patterns of acetylation stabilize the regulatory elements in an extended conformation similar to what has been observed in ex vivo patient structures. Finally, we show that a fibril with disease-associated modifications has the capacity to promote assembly in a cell model of tau aggregation. Together, our data highlight that disease-associated patterns of tau acetylation can drive tau assembly by creating new interactions not possible by a lysine residue. This study lays the groundwork for understanding how specific patterns of acetylation may mediate control of tau assembly into the various structural polymorphs seen in disease.

## Results

### Acetylation adjacent to amyloidogenic sequence regulates Tau aggregation

The frequency of acetylation on the repeat domain of tau correlates with the formation of pathological species [8], particularly when the modifications occur in proximity to amyloidogenic sequences ^306^VQIVYK^311^ and ^275^VQIINK^280^ (Fig. 1a). Using our peptide model system that leverages the regulatory elements to limit aggregation behavior^14^, we assessed whether disease-associated acetylation patterns on tau can destabilize the protective structures and drive assembly of these fragments into amyloids. We systematically probed elements that span the inter-repeat regions including R1R2, R2R3, R3R4 as well as the 3R specific isoform fragment R1R3 (Fig. 1b). The R1R2 (i.e. TENLKHQPGGGKVQI**IN**K) and R1R3 (i.e. TENLKHQPGGGKVQI**VY**K) sequences are highly homologous differing at only two positions in the amyloid motifs. Conversely, the R2R3 (i.e. **KD**N**I**KH**V**PGGG**S**VQIVYK) and R1R3 (i.e. **TE**N**L**KH**Q**PGGG**K**VQIVYK) sequences have the same amyloid motif but differ at five positions N-terminal to the amyloid motif. Our previous work on this model system uncovered that the WT sequences are aggregation resistant due to the stabilizing interactions involving the conserve P-G-G-G beta-turn stabilizing motif, and that disease-associated mutations in the R2R3 sequence drive aggregation^14^. Interestingly, testing the most dominant proline to serine mutation in the P-G-G-G motif dramatically increased R2R3 aggregation but had no effect on R1R2 and R1R3 assembly suggesting that the regulatory elements in each sequence have different capacities to reduce aggregation^14^. We hypothesize that post-translational modifications, such as acetylation, could drive these differences in aggregation.

**Figure 1.**
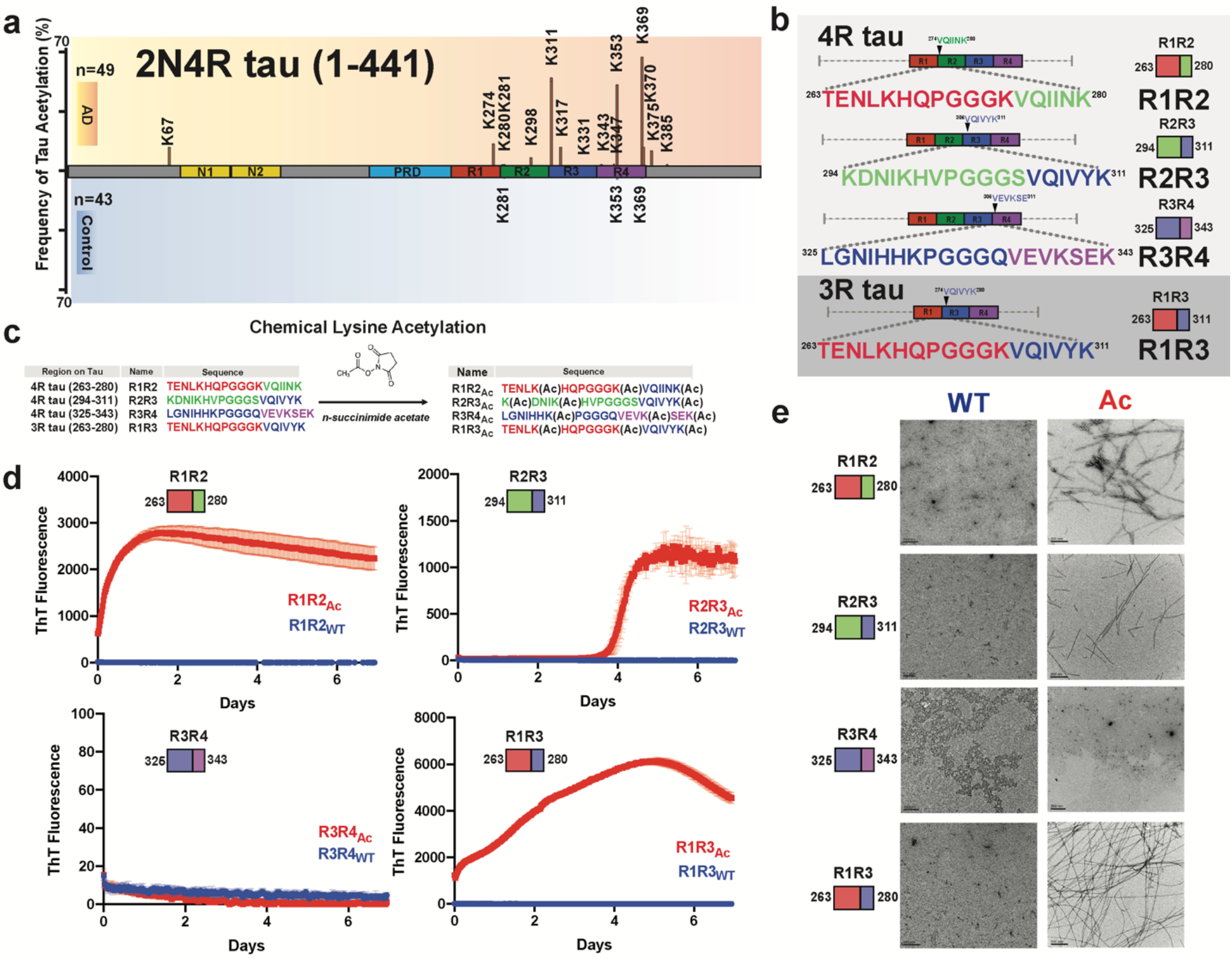
Chemical acetylation of lysines of regulatory tau fragments drives fibril assembly. Literature reported acetylation sites on 2N4R tau in AD and control patients; the modified frequency was noted on y axis. N-terminal extension domains, N1 and N2, are colored in orange. Proline-rich domain (PRD) is colored in blue. Repeat domains are colored in red, blue, green and magenta for repeats 1, 2, 3 and 4, respectively. **b.** Schematic illustrating the four tau model peptides (R1R2, R2R3, R3R4 and R1R3). Their sequence and relative location in the repeat domain are shown. Repeat domain elements are colored as in Fig. 1a. **c.** Schematic illustrating acetylation modification reaction with n-succinimide acetate of our tau model peptides. **d.** ThT fluorescence aggregation assay of acetylated peptides (red) and their corresponding non-acetylated controls (blue). Aggregation experiments were performed in triplicate and the averages are shown with standard deviation. The data were fit to a non-linear regression model fitting in GraphPad Prism to estimate an average t_1/2max_ with a standard deviation. **e.** TEM images of acetylation modified peptides (red, right column) and their corresponding non-modified controls (blue, left column). Scale bars indicate 0.2 μm.

To test the effect of acetylation on the assembly of these peptides, we utilized N-succinimidyl acetate to chemically modify lysine residues to acetyl-lysine (AcK) (Fig. 1c). Each peptide contains three lysine residues that were distributed throughout the sequence and includes lysine residues that have been observed to be acetylated in disease including K267, K274, K280 in R1R2, K311 in R2R3 and K331, K340 and K343 in R3R4 and K369 and K370^26^. All peptides contained a lysine residue at the C-terminus of the amyloid motif (i.e. ^306^VQIVYK(Ac)^311^ or ^275^VQIINK(Ac)^280^) while the other two lysines were distributed either in the N-terminus (R2R3) or one at the N-terminus and the other flanking the N-terminus of the amyloid motif (R1R2, R3R4 and R1R3). We first used mass spectrometry to confirm that the acetylation reactions proceeded to completion (Fig. 1c). We found the reaction products to be heterogenous with the most abundant species containing all three lysines acetylated but also included double and single modifications (Supplementary Fig. 1a-d). The modified peptides were purified, and their aggregation was monitored in a Thioflavin T (ThT) fluorescence aggregation assay for 7 days. Three acetylated peptides R1R2_Ac_, R1R3_Ac_, R2R3_Ac_ showed significant increase in ThT fluorescence with t_1/2max_ values of 0.20 ± 0.04, 1.66 ± 0.08 and 4.12 ± 0.01 days, respectively, while their unmodified counterparts did not aggregate (Fig. 1d). The acetylated R3R4_Ac_ and the unmodified control also did not aggregate (Fig. 1d). The presence of fibrils in the samples was confirmed with negative stain Transmission Electron Microscopy (TEM) (Fig. 1e). The R1R2_Ac_ fibrils appeared to have the most consistent twist. Our data support that chemical modification of our model tau peptides can disrupt the regulatory elements and drive their assembly into beta-sheet-rich amyloids. We next set out to identify specific patterns of acetylation on our peptides that drive the formation of fibrils.

### Defined patterns of acetylation drive tau into amyloid fibrils

With the observation that chemical acetylation of our peptides could dramatically change the aggregation propensity, we used a synthetic modification approach to generate all possible combinations of lysine acetylation modifications. R1R2, R2R3 and R1R3 peptides were synthesized with discrete modifications at each site individually (i.e. 1_Ac_, 2_Ac_, and 3_Ac_), as pairs (i.e. 12_Ac_, 23_Ac_, and 13_Ac_) and at all three sites (i.e. 123_Ac_) yielding 7 different combinations of modifications for each peptide (Fig. 2a). Additionally, we tested a subset of variants for the R3R4 and R4R’ fragments (Fig. 2b). Within this series we include acetylation modifications that have been observed in disease including K267, K274, K280 in R1R2; K311 in R2R3; K331, K340, K343 in R3R4; and K369, K370 in R4R’^26^. Aggregation of the above peptides was evaluated in a ThT aggregation assay monitored for 8 days. For both the R1R2 and R1R3 series that only vary by two positions in the amyloid motifs (Fig. 1b), we observed that modification on the second and third lysines (R1R2_23_Ac_ and R1R3_23_Ac_) or all three lysines (R1R2_123_Ac_ and R1R3_123_Ac_) aggregated rapidly with t_1/2max_ values of 0.23 ± 0.02, 1.98 ± 0.13, 0.19 ± 0.04 and 0.56 ± 0.01 days, respectively (Fig. 2c,d and Supplementary Fig. 2a). All other R1R2 and R1R3 modified peptides and their unmodified controls did not aggregate (Fig. 2c,d and Supplementary Fig. 2a). The data on the R1R2 and R1R3 peptides suggest that modification on the second and third lysines on the peptide have the largest impact the kinetics of its fibrillization. In contrast, the R2R3 series yielded much different results. While the control R2R3 peptide did not aggregate, all except for one modified peptide aggregated within the time frame of the experiment (Fig. 2e). Fitting the data to quantify the t_1/2max_ uncovers general rules in which the propensity to aggregate appear to be similarly site dependent but that the R2R3 sequence has a lower capacity to limit assembly compared to the R1R2 and R1R3 peptides. For the R2R3 peptides, we find that the 23_Ac_, 123_Ac_ and 2_Ac_ modifications yield the fastest aggregation with t_1/2max_ values of 1.8 ± 0.18, 2.56 ± 0.03 and 3.18 ± 0.31 days, respectively, suggesting that modifications on the second lysine may be the most important for mediating aggregation (Fig. 2e and Supplementary Fig 2a). The remaining R2R3 peptides formed aggregates within the time scale of the experiment with t_1/2max_ values that ranged from 3.80 to 5.35 days. The unmodified R2R3 and R2R3_3_Ac_ did not aggregate (Fig. 2e and Supplementary Fig 2a). Finally, none of the R3R4 and R4R’ peptides aggregated (Fig. 2f). For all samples, the formation of fibrils was confirmed by negative stain TEM (Fig. 2f and Supplementary Fig. 2b). Our experiments have uncovered general rules for how the regulatory tau peptides may be disrupted by acetylation sites observed in disease to drive aggregation. We find that modifications closest to the amyloid motifs have the largest effect on disruption of the protective structures.

**Figure 2.**
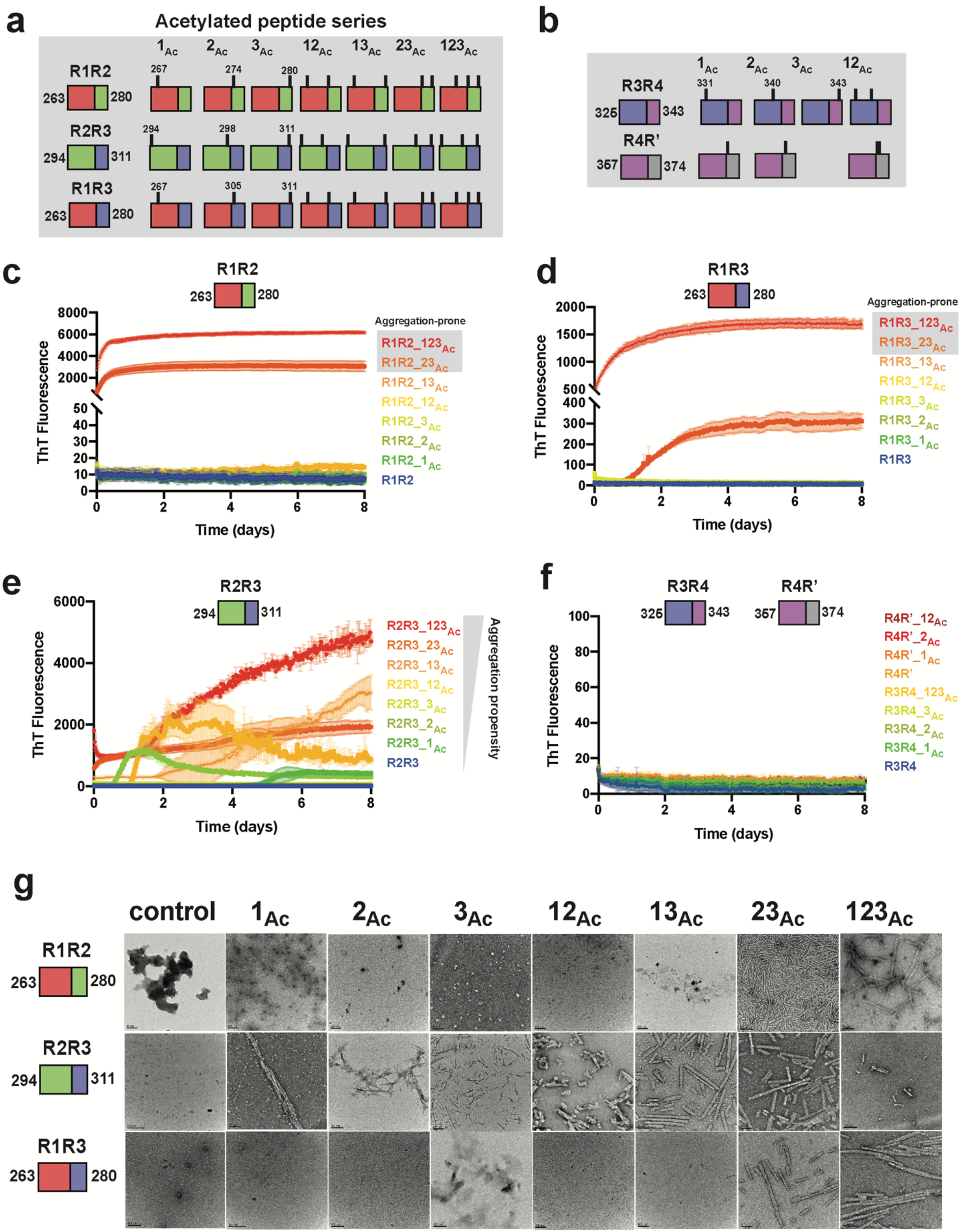
Site specific acetylation on tau peptides and their aggregation propensity. Illustration of the tau peptide series with all combinatorial acetylation patterns for the R1R2, R2R3 and R1R3 (**a**) and the R3R4 and R4R’ (**b**) tau peptides. Sequences are colored by repeat domain as in Fig. 1a. Acetylation sites are indicated by ticks above the cartoon for each peptide. ThT fluorescence aggregation experiments of the R1R2 (**c**), R2R3 (**d**), R1R3 (**e**) and R3R4/R4R’4 (**f**) unmodified and acetylation modified series. Curves are colored by the number of modifications from blue (control) to red (fully modified). Aggregation experiments were performed in triplicate and the averages are shown with standard deviation. The data were fit to a non-linear regression model fitting in GraphPad Prism to estimate an average t_1/2max_ with a standard deviation. **g**. TEM images of ThT fluorescence aggregation assay end products from control (WT) and each acetylated peptide. Scale bars indicate 0.05-0.2 μm.

### Lysine to glutamine mutations mimic acetylation in aggregation experiments

The two possible routes of lysine acetylation on proteins are possible that involve either chemical or enzymatic modification ^35^. Methods to genetically encode acetylation using orthogonal incorporation using amber codons is possible but yields and efficiency remain an issue ^36^. Genetically, the closest amino acid that mimics an acetylated lysine is glutamine but the length of the side chain and geometry of the polar groups is different^35^. Nonetheless, this is the closest approximation that has been employed in both recombinant, cellular and animal systems. Previous work on tau has shown that a lysine to glutamine substitution at position 280 promoted aggregation of full-length tau in vivo, in cells and in vitro ^30,37^, however, this modification alone in our peptide system (Fig 2c, R1R2_3_Ac_) did not aggregate. To understand the relationship between acetylation and glutamine substitutions, we generated a series of R1R2, R2R3 and R1R3 peptides with lysine to glutamine mutations at singly (i.e. 1_Q_, 2_Q_, and 3_Q_), doubly (i.e. 12_Q_, 23_Q_, and 13_Q_) and triply (i.e. 123_Q_) substituted sites (Supplementary Fig. 2c). Similar to the above experiments with acetylated peptides, the peptides were aggregated in a ThT fluorescence aggregation assay over 8 days. We found that the R1R2, R2R3 and R1R3 peptides with single modifications of lysine to glutamine on the first and second lysine positions (i.e. 1_Q_ and 2_Q_) did not aggregate with the exception of R2R3_2_Q_ which had a t_1/2max_ of 3 ± 0.3 days (Supplementary Fig. 2d-g). Glutamine substitution on the third lysine for all peptides yielded rapid aggregation that ranged from 0.16 to 1.5 days with the R2R3_3_Q_ peptide aggregating the fastest (Supplementary Fig. 2d-g). For double substitutions in the R1R2 and R1R3 sequences, the R1R3_23_Q_ mutants aggregated the fastest (t_1/2max_ < 0.25 day), followed by 12_Q_ (t_1/2max_ < 1.5 days) while the R1R3_13_Q_ mutants were the slowest (Supplementary Fig. 2d,f). The R2R3 double mutants behaved similarly except for R2R3_13_Q_ which started with fluorescence signal at time zero (Supplementary Fig. 2e). Finally, the triple substituted peptides all aggregated rapidly with t_1/2max_ values that ranged from 0.12 to 1.2 days (Supplementary Fig. 2d-f). For all incubated peptides, we confirmed the presence of fibrils using negative stain TEM (Supplementary Fig. 2h). Interestingly, the fibril morphology of the lysine to glutamine fibrils was wider than the acetylated peptides. Overall, we find that our “Q” peptide series largely follows the behavior of the acetylated peptides with perturbations on the second position being the most important for driving fibril formation. Our data also suggest that glutamine mutations appear to strongly increase aggregation propensity consistent with asparagine and glutamine residues increasing amyloidogenicity of sequences ^38,39^. More importantly, these data point to the role of positively charged lysines in slowing down assembly while polar groups of acetyl-lysine (AcK) and glutamines can promote fibril assembly. Despite being implicated in driving aggregation in disease, it has thus far remained unknown how acetylation can promote fibril assembly.

### Cryo-EM structure of an acetylated tau peptide

To gain insight into how acetylation may regulate formation of fibrils in tau, we set out to determine a cryo-EM structure of an acetylated tau fragment fibril assembly. The R1R223_Ac_ peptide aggregated rapidly and TEM imaging revealed fibrils with a defined and reproducible twist (Fig. 2d,g). We then optimized cryo-EM grids with R1R2_23_Ac_ fibrils so that the fibrils were dispersed to facilitate automatic picking of fibril filaments (Supplementary Fig. 3a). We collected 6209 images on a Titan Krios and used crYOLO to auto-pick filaments, and used Relion 3.1^40^ to reconstruct 2D and 3D maps (Supplementary Fig. 3b,c and methods). The fibril exhibited a crossover distance of 855 Å equivalent to a twisting angle of −1° and a helical raise of 4.75 Å. The fibril was assumed to have a left-handed twist since the resolution of the map, 3.9 Å, was not sufficiently high to determine the handedness unambiguously (Supplementary Fig. 3d). Each layer is comprised of 12 different fragments (Supplementary Fig. 3e,f) that fit the density well (Fig. 3c). Each fibril layer exhibits several features that are related by symmetry. The central core of the fibril is comprised of two full-length R1R2_23_Ac_ fragments (i.e. TENLKHQPGGGK(Ac)VQIINK(Ac)) that are related by a 2-fold symmetry (Supplementary Fig. 3f; i.e. A interface). This core is stabilized by a network of hydrogen bonds across two fully extended 18 residue R1R3_23_Ac_ fragments that are related by 2-fold symmetry (Supplementary Fig. 3f; i.e. A interface). Focusing on the AcK274 and Q269, we find that they form a hydrogen bond network with the 2-fold related Q269 and AcK274 of the opposite peptide fragment (Supplementary Fig. 3g). This tetravalent hydrogen bond network is the first description of a gain-of-function mediated by acetylated lysine in a cryo-EM fibril structure. Moving from this central core, the remainder of the structure is comprised of ^274^K(Ac)VQIINK(Ac)^280^ trimers that are centered on the acetylated K280 and each side contains two trimers that are related by a two-fold symmetry (Supplementary Fig. 3f,g; i.e. AcK cluster). These interactions are stabilized by contacts between a single acetyl K280 (i.e. AcK280) that forms a hydrogen bond with N279 but also nonpolar contacts to I278 and V276 (Supplementary Fig. 3g). We also find four trimeric “AcK clusters”, in which pairs of these trimers are related by a 2-fold symmetry in which two ^274^K(Ac)VQIINK(Ac)^280^ form stabilizing interactions centered on I278 (Fig. 3d and Supplementary Fig. 3g; Interface B). Similar interactions were described in x-ray crystal structures of the VQIINK amyloid motif (Supplementary Fig. 4a,b) ^41^. The periphery of these contacts is stabilized by contacts between acetyl K274 and Q276 (Supplementary Fig. 3g). In all, a single layer in our fibril structure is composed of 12 fragments that are stabilized by a combination of polar and nonpolar contacts and highlighted by the gain-of-function mediated by acetylated lysine.

**Figure 3.**
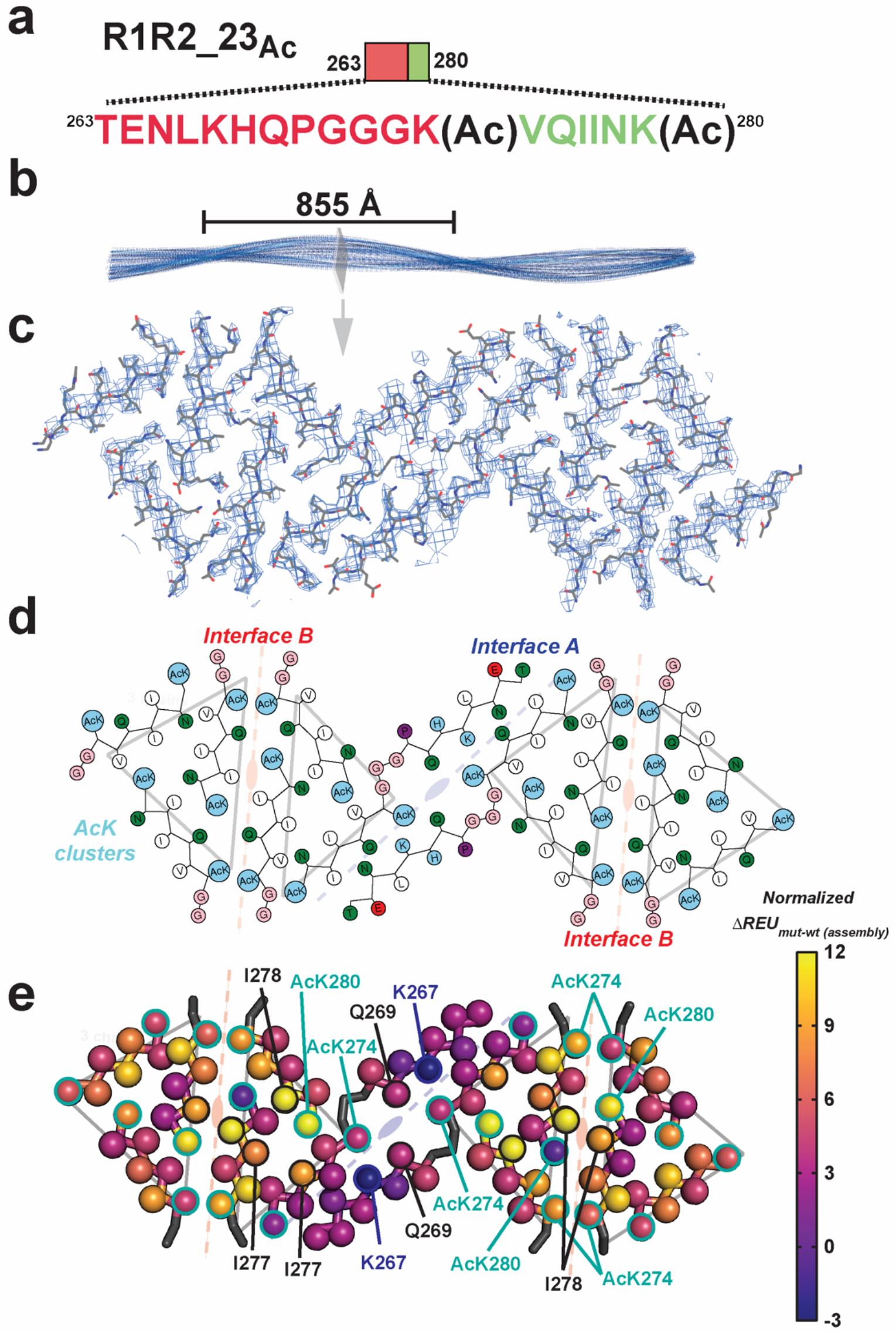
Cryo-EM structure of peptide fibril R1R2_23_Ac_. **a**. Schematic of the R1R2_23_Ac_ sequence and the acetylation sites. The sequence is colored as in Fig. 1a. Lysine acetylation sites are indicated by “Ac” following the modified lysine residue. **b**. 3D reconstruction of the fibril illustrating the cross-over distance. The fibril is shown in cartoon and is colored in blue. A single layer of the fibril is illustrated as a slice. **c**. Fit between the model and map of a single layer is contoured at 7 σ. Structure is shown as sticks with backbone and side chains. **d**. 2D illustration of the amino acid composition of the 12 molecules in the layer. AcK and lysine residues are colored light blue, nonpolar residues are white, polar residues are green, acidic residues are red and glycines are colored in pink. The description of the symmetry of the different molecules described in Fig. 3e are shown in the background. **e**. Mapping per residue 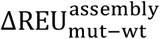 profiles onto our cryo-EM structure uncovers residues that contribute more to stability of the monolayer assembly. Structures are shown as a single layer shown in ribbons with C-beta atoms shown in spheres for each amino acid. The C-beta is colored by changes in energy due to individual amino acid mutations to alanine using the plasma color scheme, yellow (12 ΔREUs, important) to blue (−3 ΔREUs, not important). Key residues in each interface are indicated by arrows. The description of the symmetry of the different molecules described in Fig. 3e are shown in the background.

### Energetics of amino acid contributions in the R1R2_23_Ac_ fibril structure

We next interpreted the putative stability of these interactions. We leveraged a recently described computational method that captures the energetic contribution of interactions in a fibril based on changes in calculated energy of the assembly following alanine substitution^42^. Application of this method allows us to begin interpreting which interactions are important in our fibril assembly. Starting at the central “interface A”, the interactions between AcK274 and Q269 are only defined by hydrogen bonds which do not contribute significantly to the calculated energy scores (Fig. 3e and Supplementary Fig. 3g; Interface A). In this core interface, mutation of K267 to alanine is stabilizing, indicating that burial of the lysine charge may be destabilizing. Interestingly, for the R1R2 fragment, the only other acetylation modification pattern that drove assembly included acetylation on all three lysine residues, including K267 but did not yield fibrils with consistent morphologies suggesting that placement of an acetyl K267 in the context of our structure may be sterically incompatible. The other two interfaces contribute more to the stabilization. In the “interface B” site, the symmetric I278 contacts are both important for stability while the adjacent acetylated K280 contribute unevenly with one being important and the other not (Fig. 3e and Supplementary Fig. 3g; Interface B). Finally, pairs of acetylated K274’s contact each other across the interface at both ends and all four contribute modestly to the energetics (Fig. 3e and Supplementary Fig. 3g; Interface B). The “AcK cluster” interface centered on acetylated K280 which forms a hydrogen bond to N279 and nonpolar van der Waals interactions with I277 and I278 (Fig. 3e and Supplementary Fig. 3g; Ack cluster). In parallel, we used the solvation energy calculations developed by Sawaya et al ^43,44^ to assess congruence with the alanine mutational data. The calculated energetics are similar between the two methods, despite the fact that the previous method is unable to capture the contribution of acetylated lysine (Supplementary Fig. 4c,d). Thus, our data support that specific patterns of acetylation proximal to amyloid motifs may dysregulate aggregation inhibition to drive rapid fibril assembly by stabilizing new interactions not possible with lysine.

Finally, we relate our structure to previously deposited structures of tau fragments and ex vivo tauopathy assemblies. As described above, the interactions observed in the interface B of our fibril structure, resemble previously observed interactions in the x-ray crystallographic structure of KVQIINKKLD peptides (Supplementary Fig. 4a) ^41^. In the crystal structure of KVQIINKKLD peptides, V275 and I277 form 2-fold symmetric interactions (Supplementary Fig. 4a) while in a crystal structure of the minimal amyloid motif, VQIINK, I277 and I278 form stacking interactions (Supplementary Fig. 4b, i.e translational) ^41^. In our structure and within the “AcK cluster”, two I277s interact with the aliphatic chain of AcK280 while N279 makes a hydrogen bond to AcK280 (Supplementary Fig. 3g). In interface B, I278-I278 make symmetric interactions similar to VQIINK alone (Supplementary Figs. 3g and 4b) ^41^. We next compared the conformation of our structure to *ex vivo* tauopathy structures. From our structure the only region that is compatible with longer fragments of tau is the extended 18 residue fragment that forms “Interface A” (Fig. 4a). We find that the most similar element observed previously is derived from the PiD structure in which the region spanning repeats 1 and 3 similarly adopts a bulge around the P-G-G-G motif (Fig. 4b). Indeed, the alignment of the fibril core of R1R223_Ac_ and the PiD tau structure results in an root mean square deviation (rmsd) of 1.9 Å and the VQIINK segment from our structure superimposes to the VQIVYK element in the PiDs structure (Fig. 4c). Interestingly, V275 and I277 face inward to make heterotypic interactions to the ^337^VEVKSE^342^ motif in PiD while in our structure they face outward and stabilize nonpolar contacts with an acetylated lysine and I277 from a different fragment (Fig. 4d). Thus, the exposure of nonpolar residues in these elements may nucleate fibril formation as proposed previously^42^.

**Figure 4.**
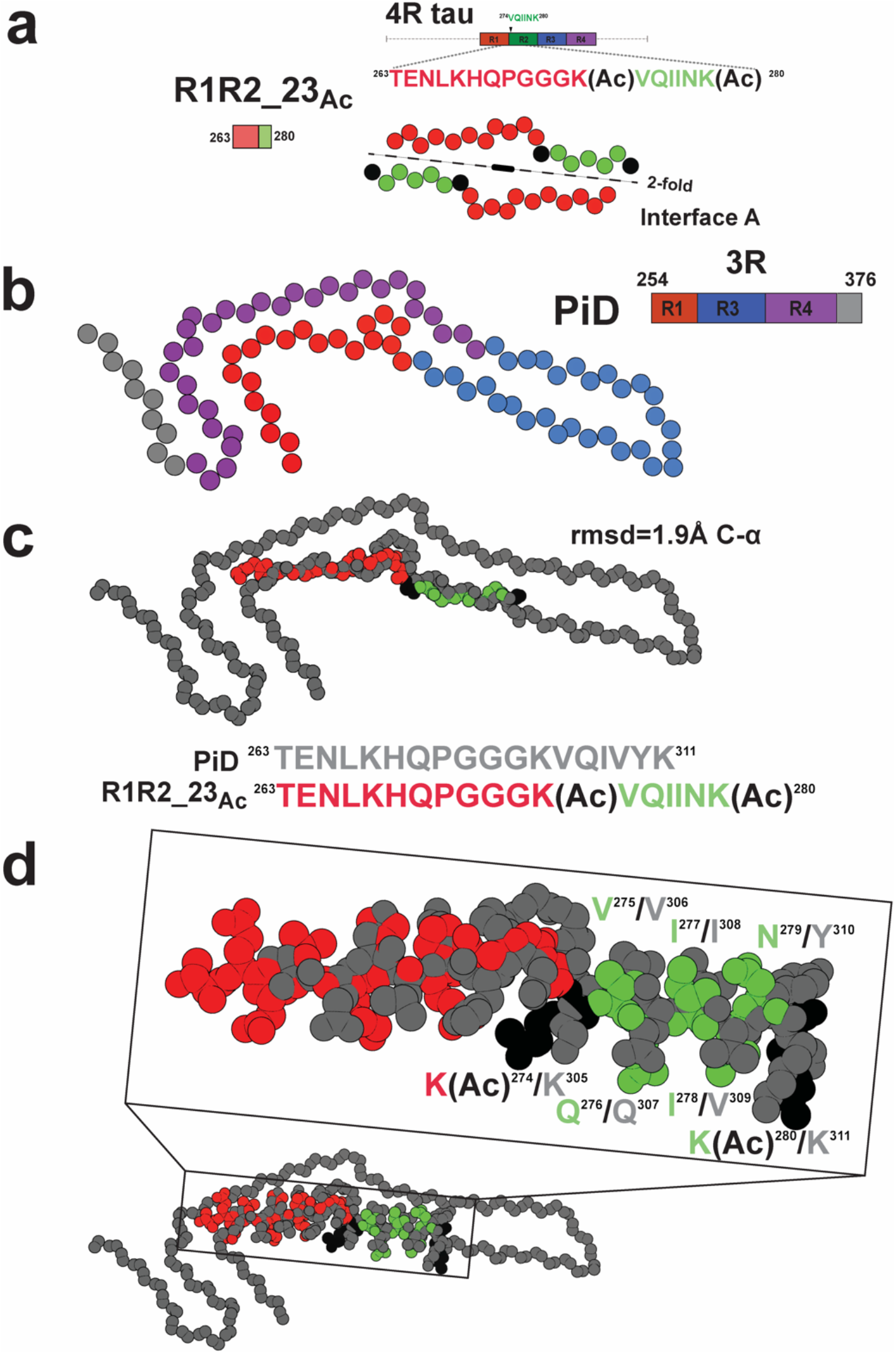
Similarity of the R1R2_23_Ac_ structure to the disease-derived Pick’s fibril. **a.** Schematic illustrating the R1R2 23_Ac_ core sequence (top). View of the core of the R1R2 23_Ac_ fragment highlighting the 2-fold “interface A” interaction (bottom). Acetylation sites are shown in black. Fragments are colored as in Fig. 1a. Structure is shown as spheres with Ca and is colored as in Fig. 1a. **b.** Illustration of a monomer layer from a protofilament of a PiD fibril (PDB id 6g×5) encoding residues 254-376. Structure and schematic is colored as in Fig. 1a. Structure is shown as spheres with C-a. **c.** Overlay of the R1R2 23_Ac_ core fragment (one chain) with the equivalent sequence in the PiD structure reveals 1.8 Å C-a atom r.m.s.d. (top). Comparison of the equivalent sequences in PiD and R1R2 23 _Ac_ that were aligned (bottom). PiD and R1R2 23_Ac_ structures/sequence are colored grey and red/green, respectively. Acetylated lysines are colored in black. The structures are shown as spheres with backbone-only. **d.** Full-atom alignment of the R1R2 23_Ac_ to the homologous region in the PiD structure. Key residues from the ^275^VQIINK^280^ and ^306^VQIVYK^311^ motifs are highlighted. Residues are numbered according to the 4R nomenclature and colored as in Fig. 4c. Structures are shown in space fill with all-atoms.

### Defined acetylation patterns on tau fragments promote seeding in cells

We also tested the biological activity of the in vitro aggregated peptides, R1R2, R2R3 and R1R3 peptides series (acetylated, glutamine mutants and WT), in a cell-based tau aggregation experiment (Fig. 5a). In this system, the tau repeat domain is expressed in HEK293T cells as fusion to FRET compatible cyan and yellow fluorescent proteins^45^. Transduction of exogenous, recombinant or patient derived, tau seeds into these tau biosensor cells converts the endogenous tau into inclusions that can be quantified by FRET^45^. For R1R2, R2R3 and R1R3 peptides, we evaluated whether *in vitro* formed aggregates containing acetyl-lysine modifications are able to seed the intracellular tau. Incubated peptides were transduced into tau biosensors and the FRET was quantified using flow cytometry to assess the extent of tau seeding (Supplementary Fig. 5a). We find that the R1R2_Ac123_ (i.e. AcK267, AcK274 and AcK280) was the only peptide in the R1R2 series that seeded significantly with 10.5±0.7% of the cells containing aggregates (Fig. 5b). In the R2R3 peptide series, the R2R3_2_Ac_ yielded signal above baseline with ~1.1% of cells containing aggregates (Fig. 5c). Unsurprisingly, the R1R3 peptides did not induce seeding consistent with there being a 4R vs 3R seeding barrier (Fig. 5d) ^46^. In parallel, we also tested activity of our “Q” peptide series and found that the R1R2_Q_ and R1R3_Q_ series showed seeding below the 1% baseline, while in the R2R3 series, R2R3_2_Q_ seeded 2.2±3.5 cells and for R2R3_123_Q_ is 1.5±0.5%. From these data, the R1R2_123_Ac_ peptide had the most seeding activity while nearly all other fragments were below 1% or slightly above this threshold (Supplementary Fig. 5b-d). Our data shows that acetylation patterns on our model peptides are important for fibril formation, however not all fibrils lead to tau aggregation in cells. Interestingly, in vitro fibril formation is not exactly linked to seeding activity in cells suggesting that other factors may be involved in propagation of seed conformation when modifications are present on the seed.

**Figure 5.**
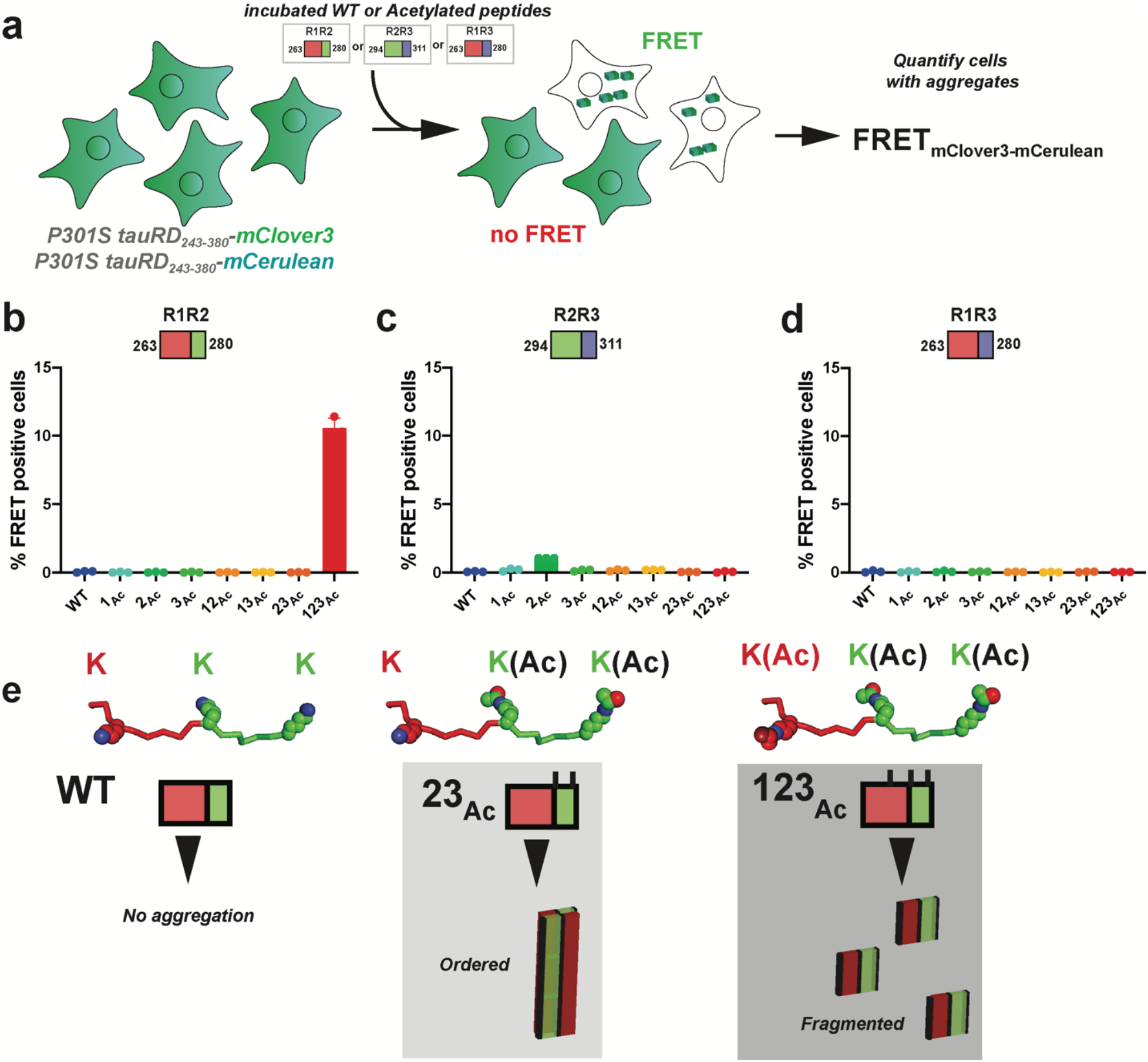
Measuring cellular tau aggregation to acetylated peptide aggregates. **a.** Schematic of the cell-based tau aggregation assay. Tau repeat domain is expressed as a fusion to mClover3 and mCerulean. Incubated peptides (WT or Acetylated) are transduced into the tau expressing cells and evaluated for conversion of the intracellular tau into aggregates as measured by FRET on a flow cytometer. Quantification of FRET signal across all peptides transduced into the tau biosensor cells: R1R2 (**b**), R2R3 (**c**) and R1R3 (**d**). Bar plots are colored as in Fig. 2. Data is shown as averages across three experiments with error bars representing a 95% CI of each condition. **e**. Model for how specific acetylation patterns on the R1R2 drive formation of fibrils but not all lead to seeding cells. The WT R1R2 sequence does not aggregate (left). The R1R2 23_Ac_ pattern (middle) leads to large ordered aggregates while the R1R2 123_Ac_ modified peptide (right) leads to more fragmented seeds which promote rapid aggregation in cells.

## Discussion

Here we present a new tau aggregation system to interpret the role of pathological acetylation patterns driving their aggregation. We identify specific acetyl lysine sites that can drive tau assembly into fibrils and show that lysine to glutamine substitutions have largely similar effects by position but lead to dramatically different fibril morphologies. We next determined a structure of a tau fibril encoding two pathological acetylation sites uncovering exciting new interactions. Consistent with our previously described tau model system, the fragments regulate tau aggregation via formation of local structures through a P-G-G-G motif and our model peptide encoding acetylation patterns uncovered an “open” conformation in the fibril of the protective hairpin. We additionally, find that acetylation on lysines promotes the formation of new interactions that balance polar and nonpolar interactions to stabilize the fibrils suggesting that the positive charge of lysine residues is inhibitory to aggregation. We also find new interactions involving the VQIINK amyloid-forming motif that interact with acetyl lysine that further stabilize the fibrils. A comparison of our new structure to pathological conformations observed in disease reveals similarities, highlighting our ability to engineer sequences that begin to control tau assembly into disease-like conformations.

### The dual roles of lysine-based regulation of tau aggregation

Under normal healthy conditions, tau is aggregation resistant and our prior work suggested that local structural rearrangements around amyloid motifs plays a role in converting tau to adopt pathological conformations even as a monomer. Using these conformational changes as a guide, we developed a tau fragment model system that highlights the role of these regulatory conformations and how pathogenic mutations disrupt them to drive aggregation. In this study we further leverage this system to show that specific acetyl lysines can also promote conversion of these protective structures into fibril assemblies. These data showcase specifically that lysine residues proximal to amyloid motifs slow down aggregation and their acetylation rapidly switches them into pathogenic conformations. Furthermore, our structure reveals that while the loss of charge on the lysine is important for fibrillization, the gain of interactions also plays a role in fibril stabilization. It is interesting to note that in all existing structures of tau, recombinant and ex vivo, lysine residues are enriched on the outside and do not stabilize the aggregates indicating that lysine residues may play protective roles. This is perhaps why tau aggregation inducers are often polyanionic, including RNA or heparin, in which the negative charges on the inducer help align the basic residues on tau to stabilize aggregation-prone conformations^10,11,13,47^. In the structures we also see that lysine residues are inhibitory to tau aggregation – evidence that acetylation can promote peptide assembly into fibrils faster. In disease, lysine residues are often expose and do not contribute to the stability of fibrils^42^ again indicating that under normal conditions they are protective but reorienting them through polyanion binding may preferentially expose the amyloid motifs to promote assembly.

### Gain of interaction of acetyl lysine groups

Classically, lysine acetylation is thought to control histone modifications that regulate DNA binding and thus exposure of sequences for subsequent transcription and gene activation^48^. It has also been proposed that enzymatic acetylation and subsequent deacetylation may modify lysines to regulate activity by destabilizing or conversely stabilizing interactions. For example, this mechanism has been proposed to regulate activity of DnaJB8 where lysine residues on the C-terminal domain may be modified to regulate its substrate activity or possibly recruitment of chaperones such as Hsp70^49^. This type of regulation has similarly been observed with phosphorylation that regulates cascades of kinases. The role of acetylation on tau remains unknown but similarly to phosphorylation, these PTMs have been proposed to increase tau’s aggregation propensity. Despite this, there are no clear patterns of PTMs for tau that drive its aggregation directly and it may simply be a modification as the protein ages or conversely a mark on the pathological assemblies following their conversion into amyloid structures. Our work has shown that tau phosphorylation may occur on tau prior to disease but there is no clear signature that marks its conversion into pathological conformations that seed^17^. Interestingly, phosphorylation on tau often occurs on the periphery outside of the repeat domain required for fibril assembly. Tau acetylation on the other hand occurs on the repeat domain but its role remains unknown. Work from Lashuel *et al* used semi-synthetic methods to produce acetylated tau at position K280 yielding modest increases in heparin-induced tau aggregation with mixed oligomer and fibril morphologies ^34^. Since lysine residues are important for tau binding to microtubules, it is possible that acetylation may influence tau function in cells and promote formation of proteotoxic species. To date, this has suggested that acetylation on tau may lead to loss-of-function with respect to microtubule binding and perhaps gain-of-function through amyloid-based proteotoxicity, but the details of these mechanisms remain unclear. Our cryo-EM structure highlights how acetylation of lysines surrounding tau amyloid motifs yields new interactions not possible with lysine residues hinting towards the formation of alternate structures with perhaps higher toxicity.

### Design of tau elements to regulate tau aggregation

Our prior work has proposed that control of amyloid motif exposure may explain tau’s capacity to assemble into distinct structural polymorphs observed in tauopathies. In a more recent analysis of tauopathy fibrils, we showed that amyloid motif interactions with other nonpolar elements in the tau sequence are essential for tau folding into different shapes. We developed ways to probe the importance of these interactions with peptides in vitro but also in cells leveraging tau’s capacity to replicate conformation in a prion-like manner. While our understanding of how these different fibrillar shapes are formed remains unknown, more general rules have emerged. We have found that the core amyloid motif interactions are modular in the ex vivo structures followed by more peripheral secondary interactions ^42^. In the present work, we demonstrate a new way dysregulate local structures and promote amyloid motif exposure using lysine acetylation, which promotes gain-of-function interactions. Interestingly, as we have shown previously the R2R3 element at the interface between repeats 2 and 3 was extremely sensitive to aggregation in response to pathogenic mutations while the R1R2 and R1R3 were aggregation resistant even in the context of the proline to serine mutations in the P-G-G-G motif. We do not yet understand how to perturb these with mutations, but our chemical modification strategy suggests that it is possible through acetylation of lysines or alternatively with mutations such as glutamine. Thus, we may be able to regulate exposure of amyloid motifs in a controlled manner to build more complex folds. Indeed, this concept was recently highlighted by Scheres *et al* where they used fragments of tau in combination with different salts to assemble structures reminiscent of ex vivo structures^50^. Thus, regulation of assembly may be further controlled by explicit exposure of amyloid motifs. Many more experiments are required to demonstrate this capacity, but the present works sets the stage to fully regulate tau assembly into conformations observed in disease.

## Conclusions

Our work supports an exciting new mechanism for how specific disease-associated lysine acetylation signatures may trigger assembly of tau towards pathogenic conformations. Structural analysis of the fibrils reveals gain of interactions mediated through lysine acetylation proximal to amyloid motifs that stabilize fibrillar conformations that partly mimic states observed in disease. Lysine residues show inhibitory effect in fibril aggregation, however modifying these lysines with acetyl groups can alter such effect to allow a mix of polar and nonpolar contacts. We envision that even though acetylation may not control final folding, it promotes interactions with amyloid motifs and perhaps plays a role in initiating tau assembly into pathological conformations. Future efforts must be focused on the design of tau elements that regulates exposure of amyloid motifs to drive formation of distinct structural polymorphs.

## ACKNOWLEDGEMENTS

L.A.J was supported by grants from the NIH-NCI (1U01CA242115), the Welch Foundation (I- 1928-20200401) and the Chan Zuckerberg Initiative (CZI) Collaborative Science Award (2018-191983). Cryo-EM data were acquired at the Cryo-Electron Microscopy Facility (CEMF) and the Structural Biology Laboratory (SBL) at UTSW, which are supported by a grant from the Cancer Prevention & Research Institute of Texas (RP170644). Transmission electron microscopy was carried out at the Electron Microscopy Core Facility at UTSW, which is supported by the National Institutes of Health (NIH) (1S10OD021685-01A1 and 1S10OD020103-01). Computational resources were provided by the BioHPC cluster supported by the Lyda Hill Department of Bioinformatics at UTSW. We would like to thank Dr. Li and Dr. Zhe Chen from the UTSW SBL core for their valuable help with cryo-EM data acquisition and analysis. We thank all members of the Joachimiak lab for discussions and input on the manuscript.

## AUTHOR CONTRIBUTIONS

L.L. and L.A.J. initiated the project. L.L. performed all peptide aggregation experiments. L.L. performed TEM of tau peptide fibrils. L.L. and B.N. determined the cryo-EM fibril structure. V.M. performed the Rosetta energy calculations. L.L. carried out the cell-based aggregation experiments. L.S. provided knowledge on cryo-EM structure determination. Finally, L.L. and L.A.J. conceived of and directed the research as well as wrote the manuscript. All authors contributed to the revisions of the manuscript.

## DATA AVAILABILITY

The structural datasets generated during the current study are available in the Electron Microscopy Data Bank repository (https://www.ebi.ac.uk/emdb/) under accession number, EMD- 28721. The structural model that was fit into the density is available in the Protein Data Bank under PDB id 8FNZ (https://doi.10.2210/pdb8FNZ/pdb). Raw ThT and cell-based aggregation data are available as source data 1 and source data 2, respectively. Any other data sets generated during and/or analyzed during the current study are available from the corresponding author on reasonable request.

## Methods

### Chemical acetylation of peptides with N-Succinimidyl Acetate

A 1 mM stock in a volume of 0.5 mL of wild type synthetic peptides R1R2, R1R3, R2R3 and R3R4 was prepared in PBS. The pH was adjusted to pH 7.4. 20 μL of 250 mM of N-succinimidyl acetate in anhydrous DMSO was added to each tube and the reactions were then incubated at room temperature RT for 30 mins. The reactions were quenched by the addition 50 μL 1 M NH4HCO3 followed by an additional 30 min incubation. The product mixtures were purified with Pierce peptide desalting spinning column (vendor) and the elutions were subsequently frozen and lyophilized (LabConco).

Both the control and acetylated peptides were reconstituted in 1mL of PBS pH 7.4 to obtain a concentration at 500μM for subsequent ThT fluorescence aggregation assays. The concentration was determined by the starting mass of peptide. In order to make an equal comparison, the WT peptides without modification were subject to the same desalting and lyophilizing procedure to minimize the concentration differences among reconstituted control and modified samples.

### Peptides with customized acetylation

Peptides with site specific acetylation were obtained from Genscript. Overall, 30 peptides were ordered with single, double and triple modifications (R1R2, R2R3, R1R3, R3R4 and R4R’, see Supplementary Table 1) including unmodified controls.

### ThT fluorescence aggregation assays

Peptides were treated with 100 μL of concentrated Trifluoroacetic acid TFA (Pierce) for 1 h at RT, in order to disaggregate any preformed aggregates. TFA was removed under nitrogen, and then lyophilized to remove any additional residue. After TFA treatment, peptides were resuspended in 1 × PBS to make 2 mM of stock, the pH was adjusted to 7.4 with NaOH. A final concentration of 500 μM of each peptide was used in 96 well plate assay with addition of 25 μM ThT (pH adjusted to 7) in a total 60 μL volume for each well. All conditions were done in triplicates. Kinetic scans were run every 30 min on a TecanSPARK plate reader at 446 nm Ex (5 nm bandwidth), 482 nm Em (5 nm bandwidth). ThT fluorescent readings of site-specific acetylated peptides and their controls were plotted in Figure 2. The t_1/2_ values were derived from a non-linear regression model fitting in GraphPad Prism. Similar method was used for K to Q mutated peptides series.

### Transmission Electron Microscopy

5 μL sample was loaded onto a glow-discharged Formvar-coated 300-mesh copper grids for 1 minute and was blotted by filter paper followed by washing the grid with 5 μL ddH_2_O. After another 30 seconds, 2% uranyl acetate was loaded on the grids for 1 minute and blotted again. The grid was dried and loaded into a FEI Tecnai G2 Spirit Biotwin TEM. All images were captured using a Gatan 2K×2K multiport readout post column CCD at the UT Southwestern EM Core Facility.

### Cryo-Electron Microscopy

The screening of cryoEM condition was performed with FEI Talos™ Arctica transmission electron microscope with K2 direct electron detector. 3 μL sample was applied to glow-discharged R1.2/1.3, 300 mesh copper grids that were plunge-frozen in liquid ethane using Vitrobot Mark IV, a Thermo Fisher Scientific Vitrobot. The optimized Vitrobot setting was: blotting force −5, blotting time 4s, humidity 98-100% at RT. Cryo-EM data were acquired at the FEI Titan™ Krios 300kV transmission electron microscope with Bioquantum energy filter, K3 direct electron detector with fringe free illustration at UT Southwestern Cryo-Electron Microscopy Facility. The prepared grids were loaded for a 24 h SerialEM run. Each grid was loaded to the stage for session set-up, grid square selection. During the fully automated SerialEM runs, each grid was loaded onto the stage, grid squares were brought to eucentric height, and holes were selected with the stored ice filter settings. All images were recorded at a dose rate of ~8 e-/pixel/s, 105,000x magnification, spot size 7, CDS mode using SerialEM software (Thermo Fisher Scientific), and converted to tiff format using relion_convert_to_tiff prior to processing. 6209 movies were collected with a 10eV energy filter slit width. Images were collected in EER mode using the aberration free image shift method in SerialEM version 3.8 (See Supplementary Table 2 for more details).

### Helical Reconstruction

Movie frames were gain-corrected, aligned and dose weighted using RELION’s motion correction program^40^. Contrast transfer function (CTF) parameters were estimated using CTFFIND-4.1 ^51^. Helical reconstructions were performed in RELION-3.1^40^. A total of 210 filaments from 24 motion-corrected movies were manually picked using EMAN2.31 ^52^ and were used to train the crYOLO 1.8.0 neural network for auto-picking ^53^. The entire dataset of 5901 movies was auto-picked and the coordinates were imported into Relion. Images of filament segments were extracted at different boxsizes of 686 and 180 pixels to perform reference free 2D classification. The best looking 2D classes of 686 pixel segments were used to estimate the cross-over distance of the filaments. The best looking 2D classes of 180 pixel segments were used to generate a de novo initial 3D reference with relion_helix_inimodel2d script ^40^ and for further 3D processing. Fibril helix is assumed to be left-handed. After multiple rounds of 3D classification, helical segments potentially leading to the best reconstructed map were chosen for 3D auto-refinements. The final 3D map was post-processed in Relion with a 6-pixel extended initial binary mask. The final overall resolution estimate was evaluated based on the FSC at 0.143 threshold between two independently refined half-maps ^54^.

### Model building and refinement

The refined map was further sharpened using phenix.auto_sharpen at the cutoff resolution of 3.88 Å ^55^. Previously published model of tau’s segments VQIINK (pdb code: 5V5C) was used as the initial template to build the filament model in COOT ^56^. Acetyl moieties were added to Lys residues by using PyTMs – a PyMOL plugin ^57^. Multiple rounds of model refinement were carried out using the default settings in phenix.real_space_refine with non-crystallographic symmetry constraints, minimization_global, rigid_body, and local_grid_search ^58^. Model geometry was evaluated using MolProbity, a built-in tool in Phenix ^59^. After each round of refinement, problematic or poorly fitting regions in the model were manually modified using COOT. This procedure was repeated until we achieved an acceptable stereochemistry model and a high overall correlation coefficients of model:map.

### *In Silico* Rosetta ΔΔG^interface^ and 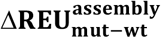 Calculation with Backrub Sampling

A four layer assembly of our cryo-EM structure was prepared using Pymol (version 2.4). Briefly, we used structural alignment to superimpose the top two chains of the deposited fibril with the bottom two chains from a duplicated fibril assembly, preserving the geometry of the assembly while extending the fibril length. Overlapping chains were removed and chains were renamed to produce assemblies of the desired number of layers with chain lettering increasing from the top to the bottom layer. To interpret energetics of acetylated lysine modified parameter files were generated for acetylated lysine. These assemblies were then used as input for the subsequent mutagenesis and minimization using the RosettaScripts interface to Rosetta^60^ in framework similar to previously described ^42^.

Changes in assembly energy were calculated using a method adapted from the Flex ddG protocol described by Barlow *et. al*.^61^. For both the 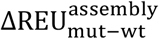 calculations, all chains were mutated. From the input assembly, a set of pairwise atom constraints with a maximum distance of 9 Å were generated with a weight of 1, using the fa_talaris2014 score function. Using this constrained score function, the structure then underwent minimization. After minimization, the residues within 8 Å of the mutation site underwent backrub sampling to better capture backbone conformational variation. These sampled structures were either only repacked and minimized, or the alanine mutation was introduced, followed by repacking and minimization. For 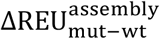 calculations, the bound wild-type and bound mutant structures reported by the interface ddG mover were used for estimating the change in assembly energy due to an alanine substitution. This is repeated for thirty-five independent replicates. The lowest energy bound mutant and bound wild-type structure energies from each replicate were extracted, and the change in energy as calculated by subtracting the wild-type, non-mutagenized assemblies’ energy from the mutant assemblies’ energy. The mean change in energy over the 35 replicates was reported as that residue’s 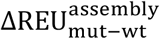.

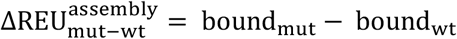

The 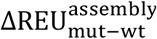 for a given residue were then averaged over 35 replicates to yield the final values for the residue. This procedure is repeated for every residue in the structure to generate a set of 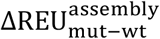 values for all residues in each fibril structure.

### Flow cytometry

The HEK193T tauRD FRET biosensor cell line developed by Diamond and colleagues can sensitively and specifically detect pathogenic tau seeds from recombinant or tissues sources ^45^. For all experiments, HEK293T tauRD biosensor cells co-expressing tauRD-mCerulean and tauRD-mClover were plated in a 96-well plate at 20,000 cells per well in 130 μL of media, and used after 24 h of plating. Peptide incubation end product (from ThT assay) were sonicated at amplitude 75 for 20 min on Q700 Sonicator 600 (QSonica). 2 μL Lipofectamine™ 2000 and 77uL Opti-MEM™ was incubated for 5 min, then 1 μL of sonicated peptide was added. The mixture was set still in cell culture hood for 20 min. 20 μL of the peptide mixture (seed mixture) was added to each well with cells. The final concentration of the peptide in each well was 0.5 μM. The seeding activity (FRET fluorescent) were visualized under fluorescent microscope after 48 h of peptide treatment, images were taken. Then the cells were harvested by 0.05% trypsin digestion and then fixed in PBS with 2% paraformaldehyde.

A BD LSRFortessa was used to quantify FRET in flow cytometry. To measure mCerulean and FRET, cells were excited with the 405 nm laser, and fluorescence was captured with a 405/50 nm and 525/50 nm filter, respectively. To measure mClover3, cells were excited with a 488 laser and fluorescence was captured with a 525/50 nm filter. To quantify FRET, we used a gating strategy where mCerulean bleed-through into the mClover3 and FRET channels was compensated using FlowJo analysis software (see Supplementary Fig. 5a for gating strategy details). As described previously, FRET signal is defined as the percentage of FRET-positive cells in all analyses. For each experiment, 10,000 cells per replicate were analyzed and each condition was analyzed in triplicate. Data analysis was performed using FlowJo v10 software (Treestar).

## Supplementary Information

Source data 1. Raw ThT experiments for R1R2, R2R3, R3R4, R1R3 and R4R’ chemically modified peptides.

Source data 2. Raw seeding data for experiments for R1R2, R2R3 and R1R3 chemically modified peptides.

## Supplementary Tables

**Supplementary Table 1.**
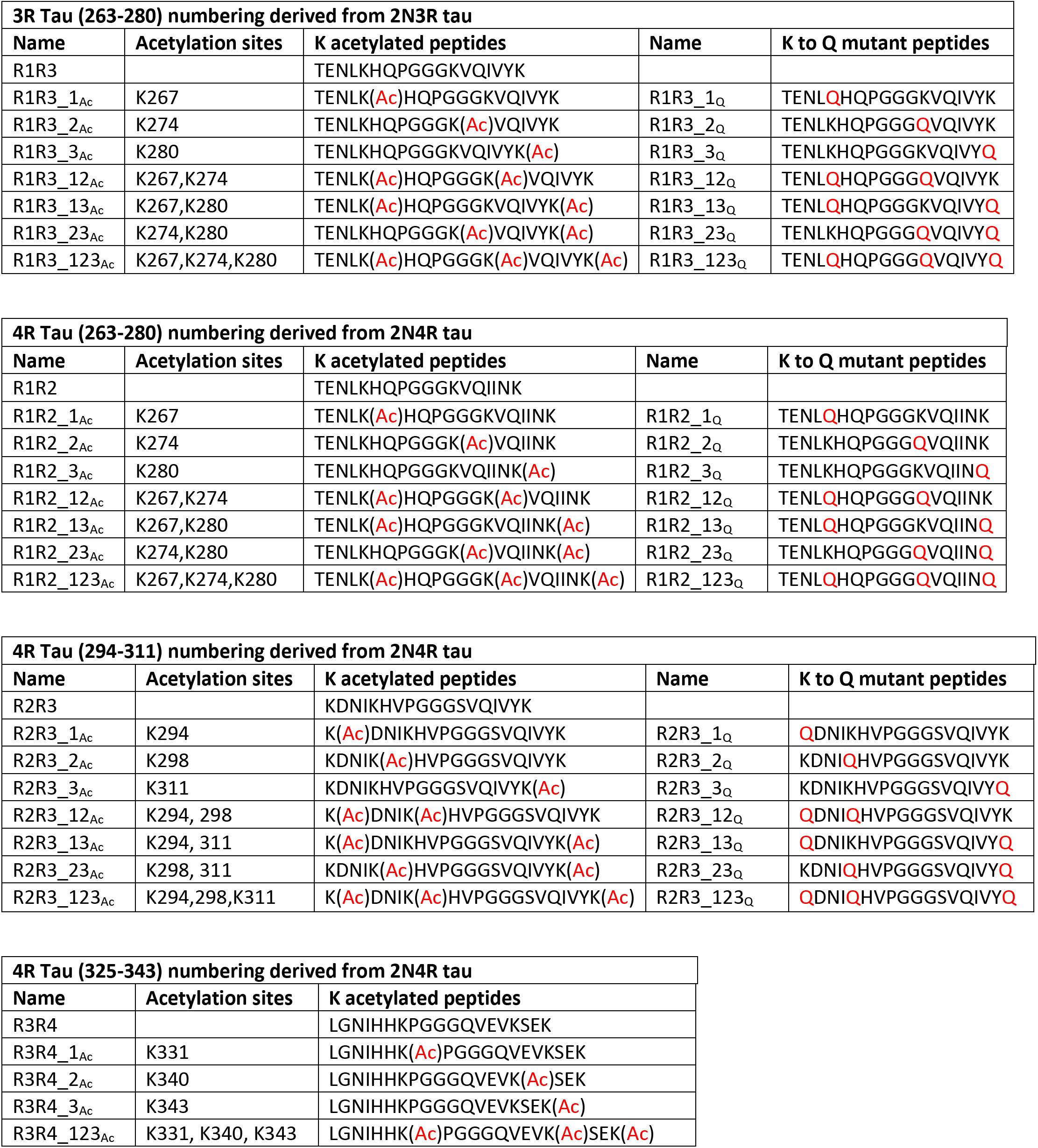
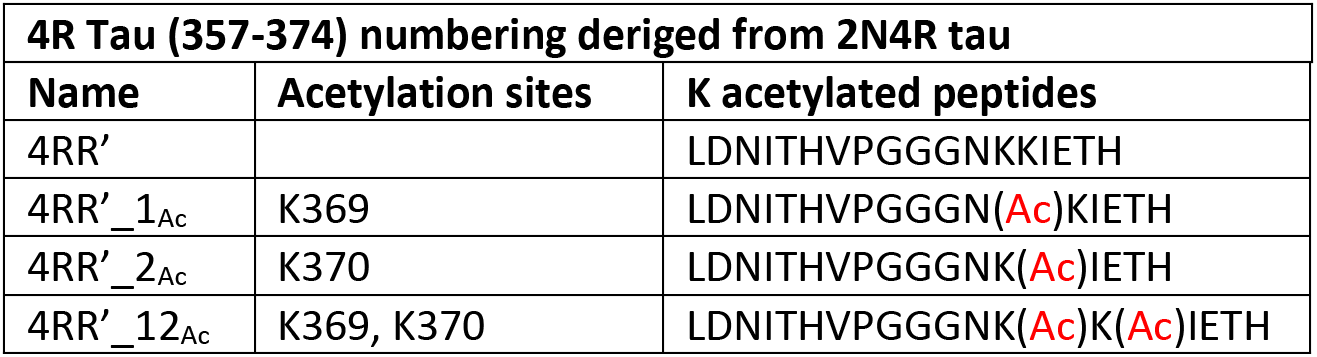
Nomenclature and sequences for all peptides used in the study. The site of acetylation or glutamine mutation is indicated by “Ac” and “Q”, respectively, and is colored in red. All peptides were N-terminally acetylated and C-terminally amidated.

**Supplementary Table 2.** Data collection and refinement statistics.

| **Data collection** |  |
| --- | --- |
| Microscope | Titan Krios |
| Acceleration Voltage (kV) | 300 |
| Detector | K3 |
| Software | SerialEM 3.8 |
| Magnification | 105,000x |
| Pixel size at detector (Å/px) | 0.83 |
| Defocus range (µm) | -1.20 to -2.40 |
| Total dose (e<sup>-</sup>/ Å<sup>2</sup>) | 52 |
| Exposure time (sec) | 4.50 |
| Number of movie frames | 50 |
| **Reconstruction** |  |
| Usable micrograph | 5901 |
| Box size (pixel) | 180 |
| Total extracted segments | 1,696,844 |
| Number of segments after 2D | 125,446 |
| Number of segments after 3D | 45,674 |
| Symmetry imposed | C1 |
| Helical rise (Å) | 4.75 |
| Helical twist (°) | -1.00 |
| Crossover length (Å) | 855 |
| Map sharpening B-factor (Å<sup>2</sup>) | -143.25 |
| Map resolution (Å; FSC=0.143) | 3.88 |
| **Model composition** |  |
| Non-hydrogen atoms | 4,096 |
| Protein residues | 504 |
| Number of chains | 4.00 |
| Water | 0.00 |
| Ligands (Acetylated) | 8.00 |
| **Model validation** |  |
| Map CC (mask) | 0.74 |
| MolProbity score | 2.14 |
| Clash score | 25.23 |
| R.M.S deviations bonds (Å) | 0.007 |
| R.M.S deviations angle (°) | 1.291 |
| Rotamer outliers (%) | 0.00 |
| Ramachandran plot<br>(favored/allowed/outliers) | (96.30/3.70/0.00) |
| Cβ outliers | 0.00 |
| CaBLAM outliers (%) | 2.56% |

## Supplementary Figures and Legends

**Supplementary Figure 1.**
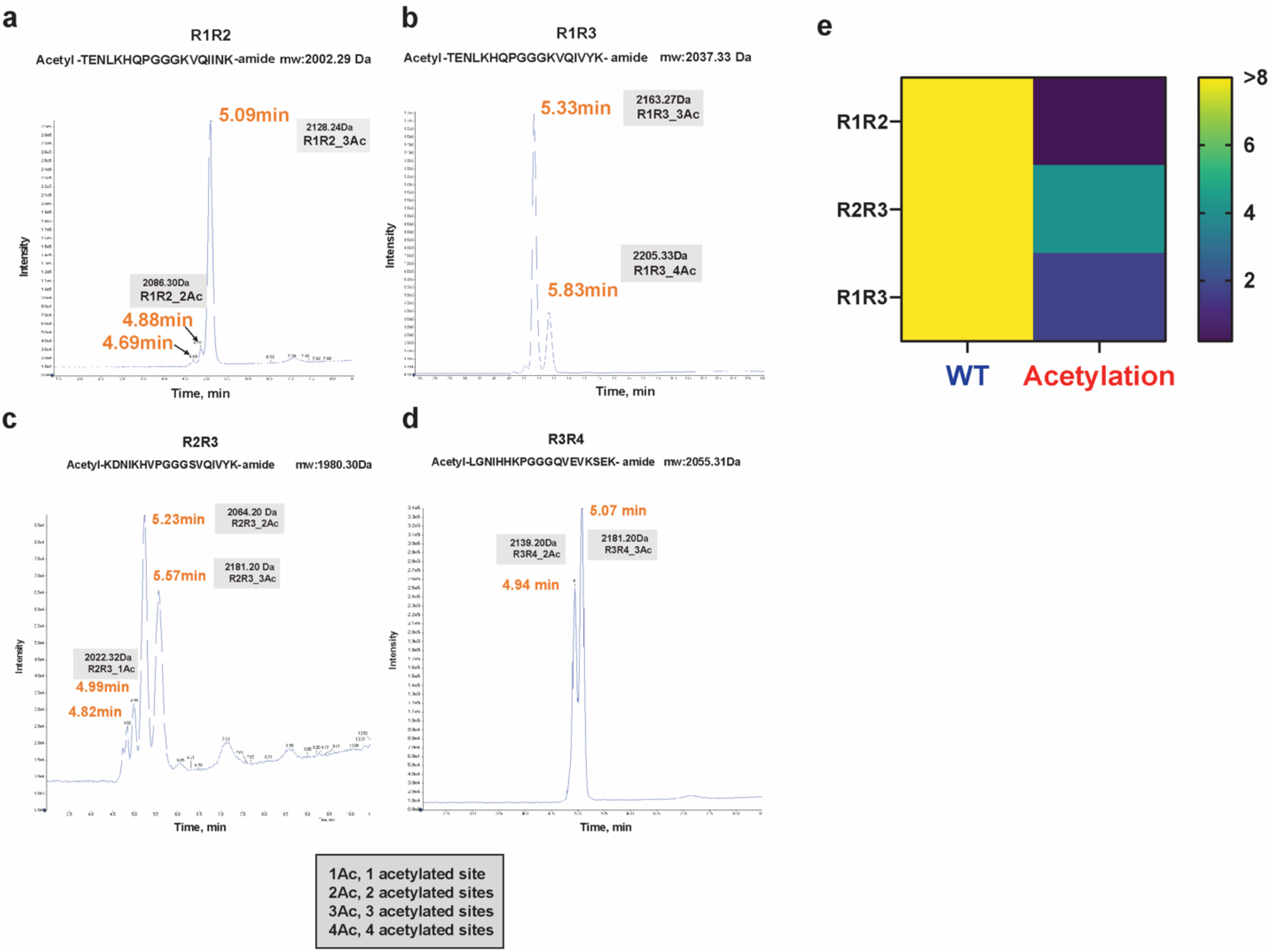
Validation of chemical modification of the R1R2, R2R3, R3R4 and R1R3 peptides. Mass spectrometry analysis of chemically acetylated R1R2 (**a**), R1R3 (**b**), R2R3 (**c**) and R3R4 (**d**) peptides. The masses for each peptide species are indicated with annotation of how many lysines are modified. **e.** Estimation of t_1/2max_ from the ThT fluorescence aggregation curves for chemically acetylated vs control peptides shown in Fig. 1d. For peptides that remained flat over the 8 days experiment we estimate the t_1/2max_ > 8 (yellow). For peptides that aggregated within 8 days, the values are colored from blue to cyan. Aggregation experiments were performed in triplicate. The data were fit to a non-linear regression model fitting in GraphPad Prism to estimate an average t_1/2max_ with a standard deviation. Error bars represent the standard deviations.

**Supplementary Figure 2.**
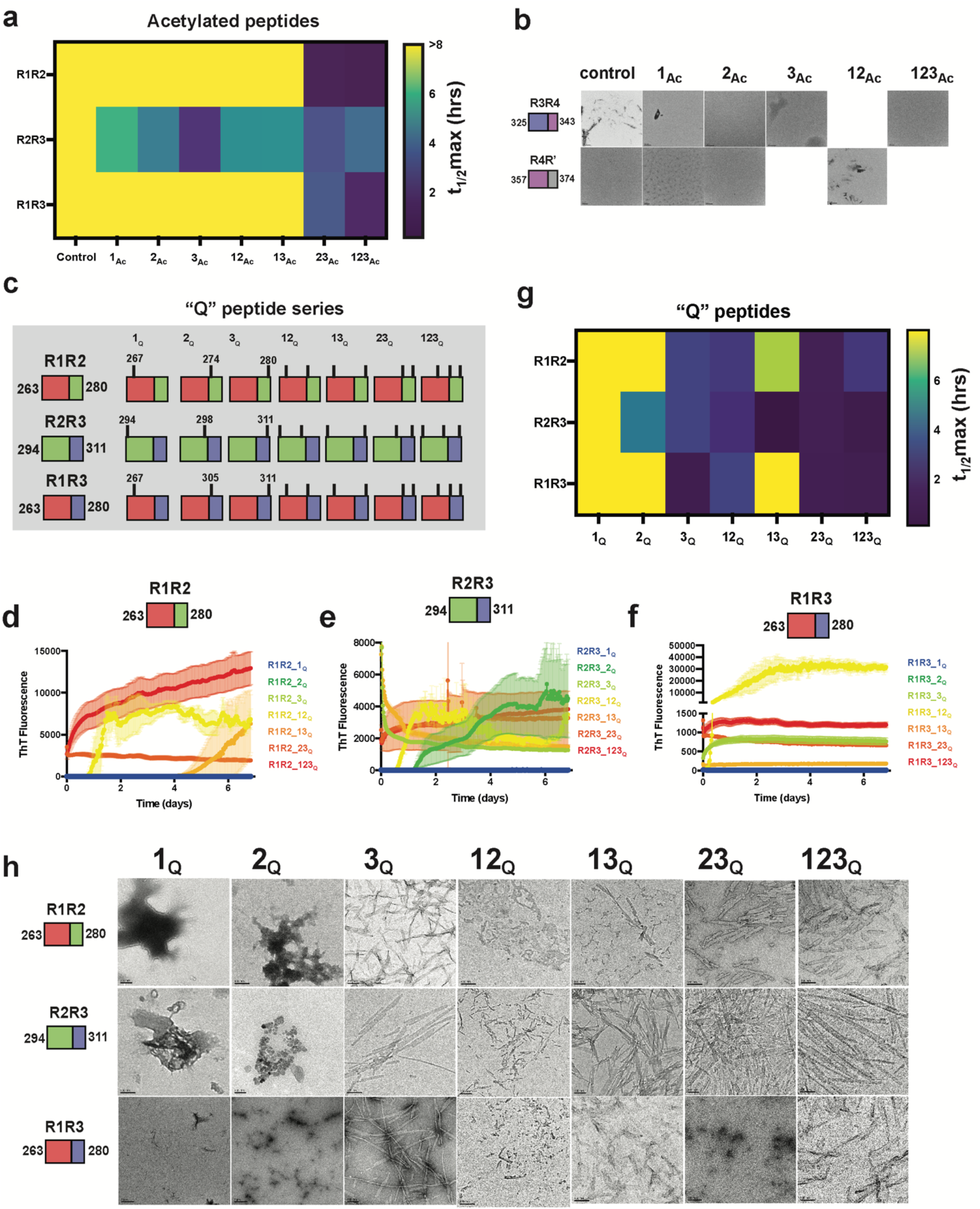
Patterns of acetylation and glutamine mutation that drive model peptide aggregation. **a**. Estimation of t_1/2max_ from the ThT fluorescence aggregation curves for unmodified, mono-, di- and tri-acetylated R1R2, R2R3, R3R4 and R1R3 peptides in Fig. 2c-f. For peptides that remained flat over the 8 days experiment we estimate the t_1/2max_ > 8 (yellow). For peptides that aggregated within 8 days, the values are colored from blue to cyan. Aggregation experiments were performed in triplicate. The data were fit to a non-linear regression model fitting in GraphPad Prism to estimate an average t_1/2max_ with a standard deviation. Error bars represent the standard deviations. **b**. TEM images of ThT fluorescence aggregation assay end products from control (WT) and the R3R4 and R4R’ acetylated peptides from aggregation experiments shown in Fig. 2f. Scale bars indicate 0.5-1μm. **c**. Illustration of the tau peptide series with all combinatorial glutamine mutations at lysine sites for the R1R2, R2R3 and R1R3 tau peptides. Sequences are colored by repeat domain as in Fig. 1a. Acetylation sites are indicated by ticks above the cartoon for each peptide. ThT fluorescence aggregation experiments of the R1R2 (**d**), R2R3 (**e**) and R1R3 (**f**) unmodified and glutamine mutated series. Curves are colored by the number of modifications from blue (control) to red (all lysines mutated to glutamine modified). Aggregation experiments were performed in triplicate and the averages are shown with standard deviation. The data were fit to a non-linear regression model fitting in GraphPad Prism to estimate an average t_1/2max_ with a standard deviation. **g**. TEM images of ThT fluorescence aggregation assay end products from control and glutamine substituted R1R2, R2R3 and R1R3 peptides from aggregation experiments shown in Supplementary Fig. 2d-f. Scale bars indicate 0.1 μm.

**Supplementary Figure 3.**
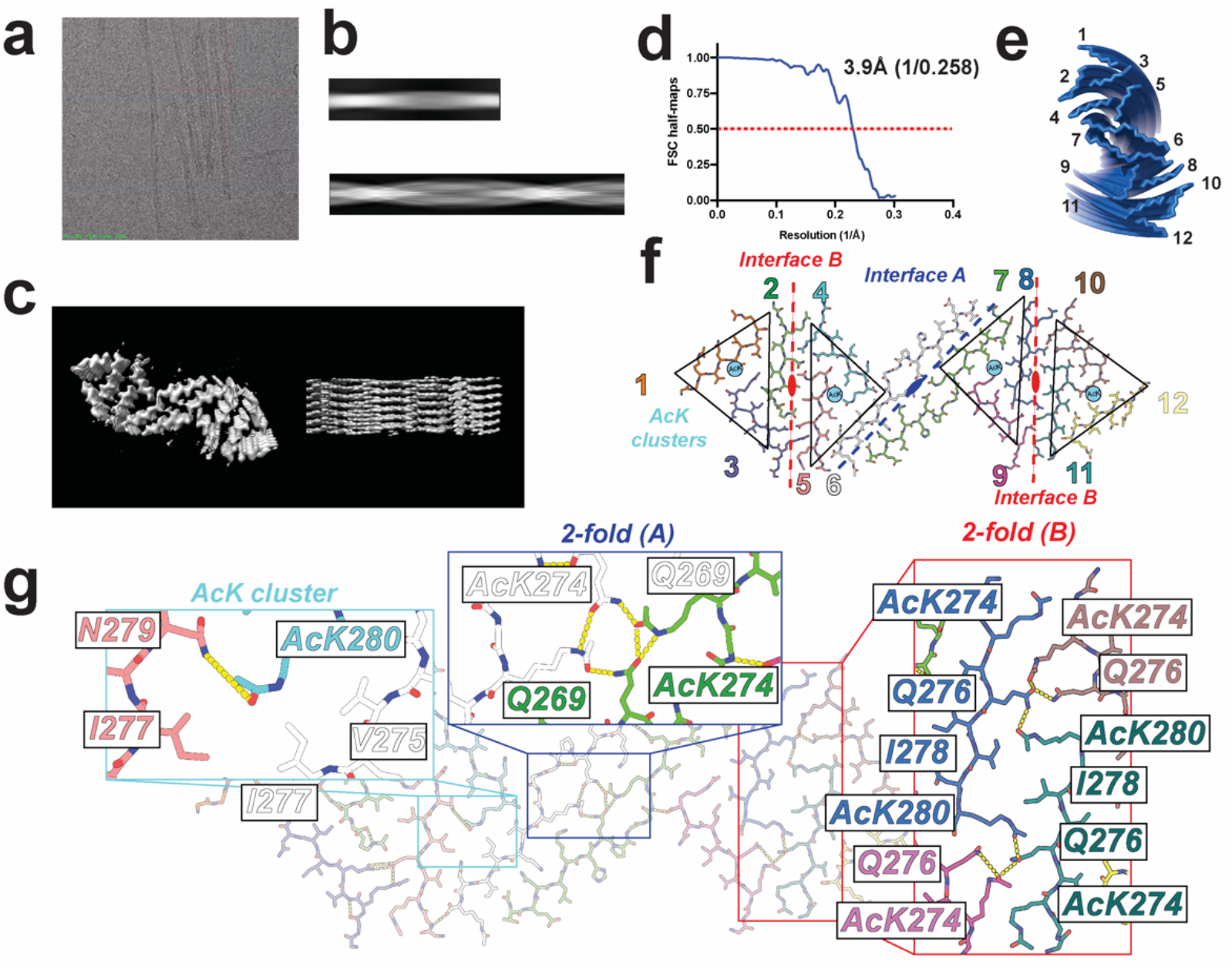
Map quality for the R1R2 23_Ac_ fibril structure. **a.** View of the R1R2 23_Ac_ fibril on a cryo-EM grid. B. 2D projection of the R1R2 23_Ac_ fibril. **c.** 3D map of the R1R2 23_Ac_ fibril contoured at 7 σ. **e.** Fourier Shell Correlation (FSC) curve for refined half-maps of R1R2 23_Ac_. **d.**Illustration of the arrangement of the 12 fragments in a single layer. **f.** Visualization of the symmetric interactions observed within a layer of our fibril assembly comprised of 12 independent chains from 1 to 12. The central interface formed between two extended fragments is annotated as “Interface A” and is defined a 2-fold symmetry axis between monomers 6-7 (blue). “Interface B” is defined by a pseudo 2-fold symmetric interactions between K(Ac)VQIINK(Ac) and K(Ac)VQIINK(Ac) from monomers 2-5 and 8-11 (red). The “interface B” interaction is flanked by a triangular interaction centered on a lysine acetylation site, termed “AcK cluster” formed between monomers 1-2-3 and 10-11-12. Structure is shown as sticks with all-atom and colored by fragment. **g.** illustration of the interactions at the different symmetric interfaces. Interface A is stabilized by three hydrogen bonds formed between two acetylated lysines at 274 and two glutamine 276 within a layer. Interface B is stabilized by symmetric interactions between K(Ac)VQIINK(Ac): K(Ac)VQIINK(Ac) mediated between I278 and I278 nonpolar contacts as as well as hydrogen bonds between acetylated K274 and Q276 from two different chains. AcK cluster is stabilized by interactions between three fragments involving nonpolar contacts between I277, I277, V275 and acetylated K280. Structure is shown as sticks with backbone and side chains.

**Supplementary Figure 4.**
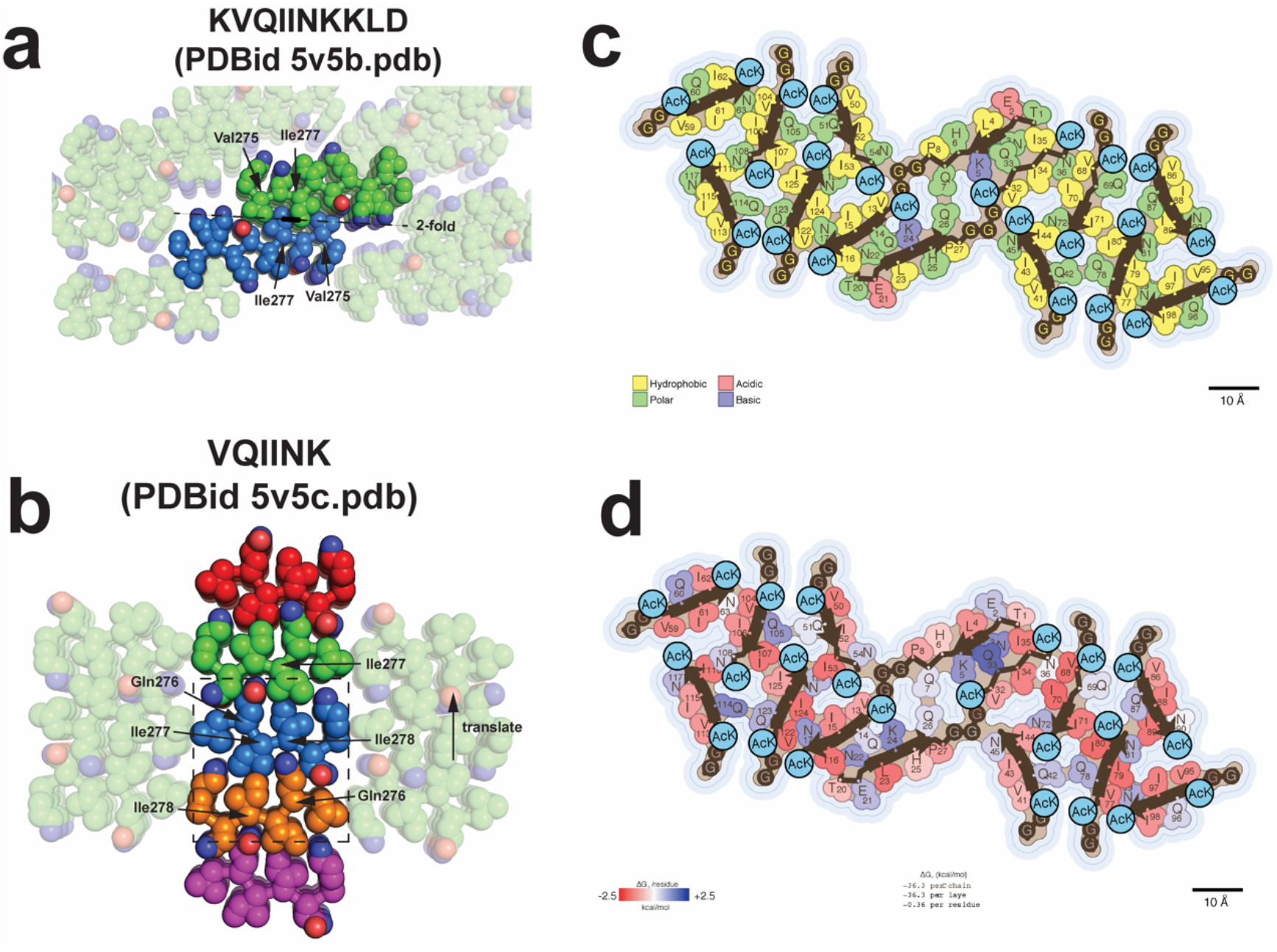
Key stabilizing interactions in R1R2 23_Ac_ are observed in structures of amyloid motif fragments. **a.** 2-fold symmetric interactions observed in X-ray structures of KVQIINKKLD (PDB id 5v5b.pdb) stabilized by interactions between V275 and I277. Structures are shown in spheres (all-atom) representation and the key interactions are highlighted. The central dimer is highlighted and the peripheral interactions are shown with transparency. **b.** X-ray structures of VQIINK (PDB id 5v5c.pdb) reveal a screw axis core dimer interaction (dashed box) that is reproduced via translation in the lattice. This interaction is stabilized by contacts between I278 and I278 with interlayer Q279 hydrogen bonding. The core of the interactions are shown as spheres (all-atoms) and colored by chain. Peripheral interactions are shown with transparency. **c**. Single layer view of the R1R2 23Ac structure highlighting the amino acid property composition across the 12 fragments. Nonpolar, polar, acidic, basic, and acetyl-lysine are colored yellow, green, red, blue and baby blue, respectively. **d**. Solvation energy estimation of per residue contribution to the stability of the monolayer. Destabilizing residues are colored in blue (+2.5 kcal/mol) and stabilizing residues are colored red (−2.5 kcal/mol).

**Supplementary Figure 5.**
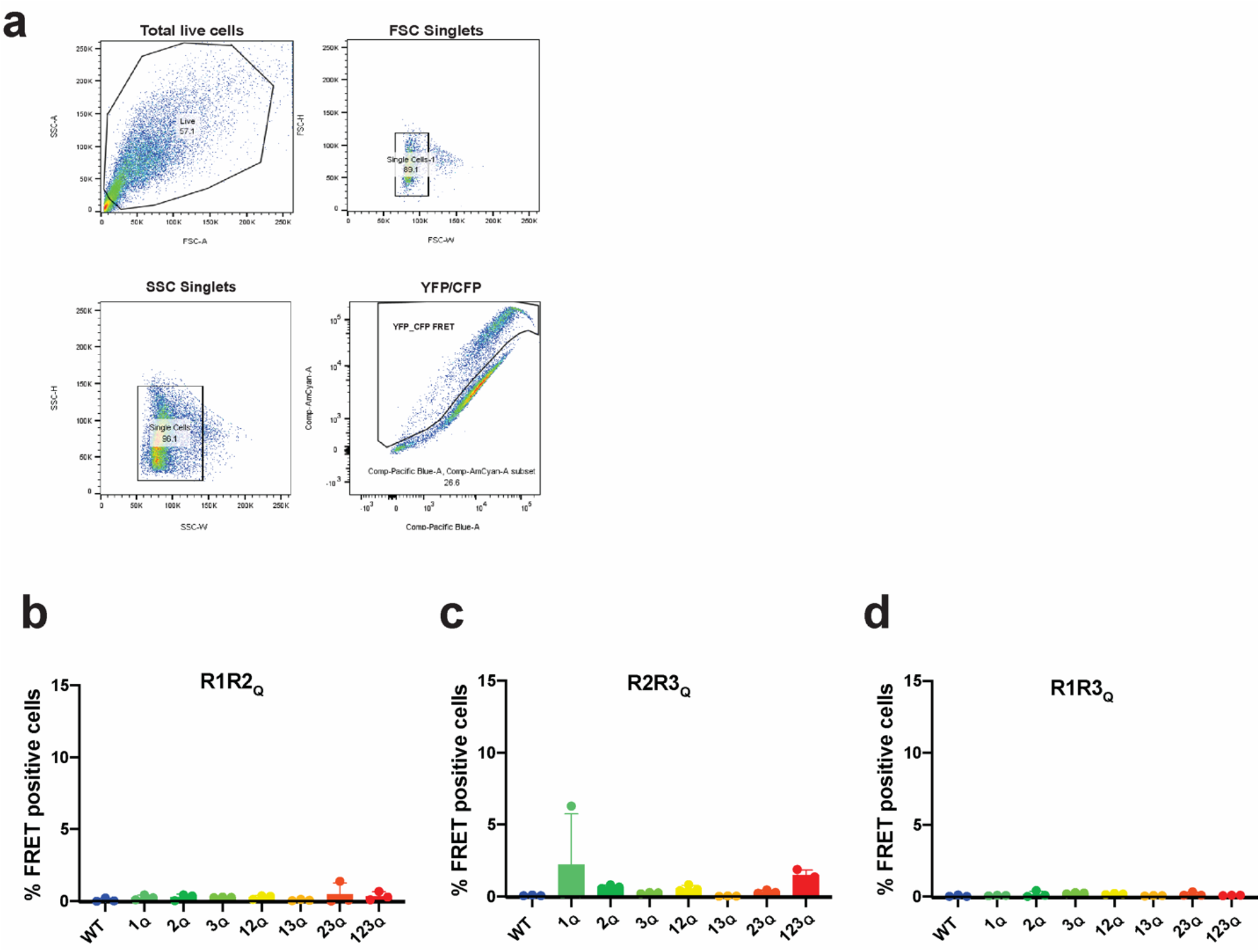
Cell-based evaluation of acetylated or glutamine substituted tau peptides. **a.** Flow cytometry strategy to quantify FRET between intracellular mClover3 and mCerulean tau aggregates. Quantification of FRET signal across control and glutamine substituted “Q” peptides transduced into the tau biosensor cells: R1R2 (**b**), R2R3 (**c**) and R1R3 (**d**). Bar plots are colored as in Supplementary Fig. 2. Data is shown as averages across three experiments with error bars representing a 95% CI of each condition.

## References

1. Vaquer-Alicea, J., Diamond, M. I. & Joachimiak, L. A. Tau strains shape disease. Acta Neuropathol. 1–15 (2021). doi:10.1007/s00401-021-02301-7

2. Fitzpatrick, A. W. P. et al. Cryo-EM structures of tau filaments from Alzheimer’s disease. Nature 547, 185–190 (2017).

3. Falcon, B. et al. Structures of filaments from Pick’s disease reveal a novel tau protein fold. Nature 561, 137–140 (2018).

4. Zhang, W. et al. Novel tau filament fold in corticobasal degeneration. Nature 580, 283–287 (2020).

5. Falcon, B. et al. Novel tau filament fold in chronic traumatic encephalopathy encloses hydrophobic molecules. Nature 568, 420–423 (2019).

6. Sanders, D. W. et al. Distinct Tau Prion Strains Propagate in Cells and Mice and Define Different Tauopathies. Neuron 82, 1271–1288 (2014).

7. Kaufman, S. K. et al. Tau Prion Strains Dictate Patterns of Cell Pathology, Progression Rate, and Regional Vulnerability In Vivo. Neuron 1–17 (2016). doi:10.1016/j.neuron.2016.09.055

8. Clavaguera, F. et al. Brain homogenates from human tauopathies induce tau inclusions in mouse brain. Proc. Natl. Acad. Sci. U.S.A. 110, 9535–9540 (2013).

9. Guo, J. L. & Lee, V. M.-Y. Cell-to-cell transmission of pathogenic proteins in neurodegenerative diseases. Nat. Med. 20, 130–138 (2014).

10. Goedert, M. et al. Assembly of microtubule-associated protein tau into Alzheimer-like filaments induced by sulphated glycosaminoglycans. Nature 383, 550–553 (1996).

11. Zwierzchowski-Zarate, A. N. et al. RNA induces unique tau strains and stabilizes Alzheimer’s disease seeds. J. Biol. Chem. 298, 102132 (2022).

12. Hou, Z., Chen, D., Ryder, B. D. & Joachimiak, L. A. Biophysical properties of a tau seed. Nature Publishing Group 11, 13602–9 (2021).

13. Mirbaha, H. et al. Inert and seed-competent tau monomers suggest structural origins of aggregation. eLife 7, 338 (2018).

14. Chen, D. et al. Tau local structure shields an amyloid-forming motif and controls aggregation propensity. Nature Communications 10, 2493–14 (2019).

15. Mukrasch, M. D. et al. Highly populated turn conformations in natively unfolded tau protein identified from residual dipolar couplings and molecular simulation. J. Am. Chem. Soc. 129, 5235–5243 (2007).

16. Stelzl, L. S. et al. Global Structure of the Intrinsically Disordered Protein Tau Emerges from its Local Structure. bioRxiv 2021.11.23.469691 (2022). doi:10.1101/2021.11.23.469691

17. Mirbaha, H. et al. Seed-competent tau monomer initiates pathology in a tauopathy mouse model. J. Biol. Chem. 102163 (2022). doi:10.1016/j.jbc.2022.102163

18. Kinoshita, J. & Clark, T. Alzforum. Methods Mol. Biol. 401, 365–381 (2007).

19. Bancher, C. et al. Accumulation of abnormally phosphorylated tau precedes the formation of neurofibrillary tangles in Alzheimer’s disease. Brain Research 477, 90–99 (1989).

20. Alonso, A. C., Grundke-Iqbal, I. & Iqbal, K. Alzheimer’s disease hyperphosphorylated tau sequesters normal tau into tangles of filaments and disassembles microtubules. Nat. Med. 2, 783–787 (1996).

21. Schwalbe, M. et al. Structural Impact of Tau Phosphorylation at Threonine 231. Structure 23, 1448–1458 (2015).

22. Despres, C. et al. Identification of the Tau phosphorylation pattern that drives its aggregation. Proc. Natl. Acad. Sci. U.S.A. 114, 9080–9085 (2017).

23. Barthélemy, N. R. et al. A soluble phosphorylated tau signature links tau, amyloid and the evolution of stages of dominantly inherited Alzheimer’s disease. Nat. Med. 26, 398–407 (2020).

24. Hyman, B. T. et al. National Institute on Aging-Alzheimer’s Association guidelines for the neuropathologic assessment of Alzheimer’s disease. Alzheimers Dement 8, 1–13 (2012).

25. Braak, H., Alafuzoff, I., Arzberger, T., Kretzschmar, H. & Del Tredici, K. Staging of Alzheimer disease-associated neurofibrillary pathology using paraffin sections and immunocytochemistry. Acta Neuropathol. 112, 389–404 (2006).

26. Wesseling, H. et al. Tau PTM Profiles Identify Patient Heterogeneity and Stages of Alzheimer’s Disease. Cell 183, 1699–1713.e13 (2020).

27. Haj-Yahya, M. et al. Site-Specific Hyperphosphorylation Inhibits, Rather than Promotes, Tau Fibrillization, Seeding Capacity, and Its Microtubule Binding. Angew. Chem. Int. Ed. Engl. 59, 4059–4067 (2020).

28. Arakhamia, T. et al. Posttranslational Modifications Mediate the Structural Diversity of Tauopathy Strains. Cell 180, 633–644.e12 (2020).

29. Irwin, D. J. et al. Acetylated tau, a novel pathological signature in Alzheimer’s disease and other tauopathies. Brain 135, 807–818 (2012).

30. Trzeciakiewicz, H. et al. A Dual Pathogenic Mechanism Links Tau Acetylation to Sporadic Tauopathy. Nature Publishing Group 7, 44102–13 (2017).

31. Cohen, T. J. et al. The acetylation of tau inhibits its function and promotes pathological tau aggregation. Nature Communications 2, 252–9 (2011).

32. Brotzakis, Z. F. et al. A Structural Ensemble of a Tau-Microtubule Complex Reveals Regulatory Tau Phosphorylation and Acetylation Mechanisms. ACS Cent Sci 7, 1986–1995 (2021).

33. Caballero, B. et al. Acetylated tau inhibits chaperone-mediated autophagy and promotes tau pathology propagation in mice. Nature Communications 12, 2238–18 (2021).

34. Haj-Yahya, M. & Lashuel, H. A. Protein Semisynthesis Provides Access to Tau Disease-Associated Post-translational Modifications (PTMs) and Paves the Way to Deciphering the Tau PTM Code in Health and Diseased States. J. Am. Chem. Soc. 140, 6611–6621 (2018).

35. Neumann-Staubitz, P., Lammers, M. & Neumann, H. Genetic Code Expansion Tools to Study Lysine Acylation. Adv Biol (Weinh) 5, e2100926 (2021).

36. Neumann, H. et al. A method for genetically installing site-specific acetylation in recombinant histones defines the effects of H3 K56 acetylation. Mol Cell 36, 153–163 (2009).

37. Gorsky, M. K., Burnouf, S., Dols, J., Mandelkow, E. & Partridge, L. Acetylation mimic of lysine 280 exacerbates human Tau neurotoxicity in vivo. Nature Publishing Group 6, 22685–12 (2016).

38. Halfmann, R. et al. Opposing Effects of Glutamine and Asparagine Govern Prion Formation by Intrinsically Disordered Proteins. Molecular Cell 43, 72–84 (2011).

39. Shattuck, J. E., Waechter, A. C. & Ross, E. D. The effects of glutamine/asparagine content on aggregation and heterologous prion induction by yeast prion-like domains. prion 11, 249–264 (2017).

40. Scheres, S. H. W. Amyloid structure determination in RELION-3.1. Acta Crystallogr D Struct Biol 76, 94–101 (2020).

41. Seidler, P. M. et al. Structure-based inhibitors of tau aggregation. Nat Chem 10, 170–176 (2018).

42. Mullapudi, V. et al. Network of hotspot interactions cluster tau amyloid folds. bioRxiv 2022.07.01.498342 (2022). doi:10.1101/2022.07.01.498342

43. Sawaya, M. R., Hughes, M. P., Rodriguez, J. A., Riek, R. & Eisenberg, D. S. The expanding amyloid family: Structure, stability, function, and pathogenesis. Cell 184, 4857–4873 (2021).

44. Eisenberg, D. & McLachlan, A. D. Solvation energy in protein folding and binding. Nature 319, 199–203 (1986).

45. Holmes, B. B. et al. Proteopathic tau seeding predicts tauopathy in vivo. doi:10.1073/pnas.1411649111

46. Schoch, K. M. M. et al. Increased 4R-Tau Induces Pathological Changes in a Human-Tau Mouse Model. Neuron 90, 941–947 (2016).

47. Hou, Z., Chen, D., Ryder, B. D. & Joachimiak, L. A. Biophysical properties of a tau seed. bioRxiv 2021.03.30.437772 (2021). doi:10.1101/2021.03.30.437772

48. Gräff, J. & Tsai, L.-H. Histone acetylation: molecular mnemonics on the chromatin. Nat Rev Neurosci 14, 97–111 (2013).

49. Hageman, J. et al. A DNAJB chaperone subfamily with HDAC-dependent activities suppresses toxic protein aggregation. Mol Cell 37, 355–369 (2010).

50. Lövestam, S. et al. Assembly of recombinant tau into filaments identical to those of Alzheimer’s disease and chronic traumatic encephalopathy. eLife 11, (2022).

51. Rohou, A. & Grigorieff, N. CTFFIND4: Fast and accurate defocus estimation from electron micrographs. Journal of Structural Biology 192, 216–221 (2015).

52. Tang, G. et al. EMAN2: an extensible image processing suite for electron microscopy. Journal of Structural Biology 157, 38–46 (2007).

53. Wagner, T. et al. Two particle-picking procedures for filamentous proteins: SPHIRE-crYOLO filament mode and SPHIRE-STRIPER. Acta Crystallogr D Struct Biol 76, 613–620 (2020).

54. Chen, S. et al. High-resolution noise substitution to measure overfitting and validate resolution in 3D structure determination by single particle electron cryomicroscopy. Ultramicroscopy 135, 24–35 (2013).

55. Terwilliger, T. C., Sobolev, O. V., Afonine, P. V. & Adams, P. D. Automated map sharpening by maximization of detail and connectivity. Acta Crystallogr D Struct Biol 74, 545–559 (2018).

56. Emsley, P., Lohkamp, B., Scott, W. G. & Cowtan, K. Features and development of Coot. Acta Crystallogr D Biol Crystallogr 66, 486–501 (2010).

57. Warnecke, A., Sandalova, T., Achour, A. & Harris, R. A. PyTMs: a useful PyMOL plugin for modeling common post-translational modifications. BMC Bioinformatics 15, 370–12 (2014).

58. Afonine, P. V. et al. Real-space refinement in PHENIX for cryo-EM and crystallography. Acta Crystallogr D Struct Biol 74, 531–544 (2018).

59. Chen, V. B. et al. MolProbity: all-atom structure validation for macromolecular crystallography. Acta Crystallogr D Biol Crystallogr 66, 12–21 (2010).

60. Fleishman, S. J. et al. RosettaScripts: a scripting language interface to the Rosetta macromolecular modeling suite. PLoS ONE 6, e20161 (2011).

61. Barlow, K. A. et al. Flex ddG: Rosetta Ensemble-Based Estimation of Changes in Protein-Protein Binding Affinity upon Mutation. J Phys Chem B 122, 5389–5399 (2018).

